# A Bright, Photostable Dye that Enables Multicolor, Time Lapse, and Super-Resolution Imaging of Acidic Organelles

**DOI:** 10.1101/2023.08.04.552058

**Authors:** Lauren Lesiak, Neville Dadina, Shuai Zheng, Marianne Schelvis, Alanna Schepartz

**Author notes:** **Corresponding Author Alanna Schepartz** – Department of Molecular and Cell Biology and Department of Chemistry, University of California, Berkeley, California 94720, United States; Molecular Biophysics and Integrated Bioimaging Division, Lawrence Berkeley National Laboratory, Berkeley, California 94720, United States; Chan Zuckerberg Biohub, San Francisco, California 94158, United States.

## Abstract

Lysosomes have long been known for their acidic lumen and efficient degradation of cellular byproducts. In recent years it has become clear that their function is far more sophisticated, involving multiple cell signaling pathways and interactions with other organelles. Unfortunately, their acidic interior, fast dynamics, and small size makes lysosomes difficult to image with fluorescence microscopy. Here we report a far-red small molecule, HMSiR_680_-Me, that fluoresces only under acidic conditions, causing selective labeling of acidic organelles in live cells. HMSiR_680_-Me can be used alongside other far-red dyes in multicolor imaging experiments and is superior to existing lysosome probes in terms of photostability and maintaining cell health and lysosome motility. We demonstrate that HMSiR_680_-Me is compatible with overnight time lapse experiments, as well as time lapse super-resolution microscopy with a fast frame rate for at least 1000 frames. HMSiR_680_-Me can also be used alongside silicon rhodamine dyes in a multiplexed super-resolution microscopy experiment to visualize interactions between the inner mitochondrial membrane and lysosomes with only a single excitation laser and simultaneous depletion. We envision this dye permitting more detailed study of the role of lysosomes in dynamic cellular processes and disease.

## INTRODUCTION

Lysosomes are essential for cellular function. As the cell’s degradative signaling hub, lysosomes consume macromolecules delivered to them by endosomes and autophagosomes and use the products to provide other organelles with building blocks for metabolism and information for cell signaling.^1^ Their ability to degrade luminal proteins has also been leveraged recently as a strategy to eliminate proteins that cause disease.^2,3^ All of these degradation events are enabled by protein or lipid hydrolases that function at low pH, and as a result the luminal interiors of lysosomes are necessarily acidic. Dysfunction of these hydrolases contributes to multiple lysosomal storage disorders including Niemann-Pick and Pompe diseases, as well as diseases associated with aging, cancer, and neurodegeneration.^4-6^

An improved understanding of lysosomes in cell function and disease would be enabled by tools to visualize their activity and dynamics in molecular detail. Unfortunately, the very nature of lysosomal function makes them exceptionally challenging to image. Their highly acidic lumen degrades many molecules used as fluorescent probes, including proteins and small molecules.^7^ Furthermore, capturing dynamic interactions with other organelles demands an imaging modality that supports a high frame rate. As a result, the fluorescent probe used must be sufficiently bright to compensate for the dim signal that results from a low exposure time per frame.^8^ Complicating matters further is the fact that individual lysosomes can be difficult to resolve, as their median size (∼400 nm) is close to the diffraction limit.^9^ Super-resolution microscopy (SRM) is a promising technique for imaging small vesicles, but it requires SRM-compatible dyes that emit in the far-red and are also bright and photostable.^10,11^ While other subcellular organelles have been thoroughly investigated,^10,12-14^ the low pH and small size of lysosomes makes studying their dynamics a major challenge.

Current imaging techniques have not yet been able to overcome these aforementioned challenges associated with studying acidic organelles. Fluorescent proteins can be conveniently fused to organelle-resident proteins to localize fluorescent signals, but they lack the brightness and photostability required to support the acquisition of high-resolution images using SRM.^7^ Protein tags that covalently bind a small-molecule fluorophore provide more spectral options and better photostability, but still require transfection of the cells under study and thus cannot be used in many non-model cell lines.^15-17^ Small-molecule fluorophores are well suited to address these problems, as they are compatible with virtually any cell type and the labeling protocol requires only a simple same-day incubation. Indeed, commercially available LysoTracker™ dyes and recently reported LysoPB Yellow, among others, share these benefits.^18-26^ However, only a handful of reported lysosome probes, including LysoTracker Deep Red (LTDR), absorb above 600 nm, which is required for long-term live-cell imaging as higher energy excitation is cytotoxic. Unfortunately, these dyes lack the photostability required for long-time lapse SRM.^27-30^

The ideal candidate for imaging lysosomes would be a bright and photostable small molecule that absorbs above 600 nm and labels lysosomes without need for a specific targeting moiety. Rhodamines are bright and photostable small molecules and their modular structure allows accurate fine-tuning of spectral and chemical properties.^11,31-33^ Silicon rhodamines, in which oxygen is replaced with dimethyl silicon, absorb above 600 nm and have been widely utilized for SRM.^34-36^ What is missing is a transfection-free way to target a silicon rhodamine fluorophore to the lysosome. Here we report the design and application of HMSiR_680_-Me, a bright, photostable silicon rhodamine fluorophore that absorbs at 680 nm and fluoresces only in acidic environments. HMSiR_680_-Me enables lysosomes to be imaged in live cells for extended times, at super resolution, and in multiple colors.

## RESULTS AND DISCUSSION

When we set out to design a SRM-compatible lysosome probe, we were particularly intrigued by the photophysical properties of HMSiR_THQ_, a recently reported far-red silicon rhodamine fluorophore whose emission intensity showed a pH dependence.^36^ Specifically, HMSiR_THQ_ is non-fluorescent at physiological pH (pH 7.4) but becomes highly fluorescent at pH values below 6. This pH dependence is a direct result of an intramolecular cyclization reaction that interconverts a non-fluorescent cyclized form of the dye, which predominates at high pH, with a fluorescent, open form that predominates at lower pH (Figure 1a). Notably, the fluorescent form that predominates at lower pH benefits from both a reasonable quantum yield (0.38) and an emission maximum of almost 700 nm. The midpoint of this pH-dependent equilibrium (p*K*_cycl_) is 6.9, just below cytosolic pH.

**Figure 1.**
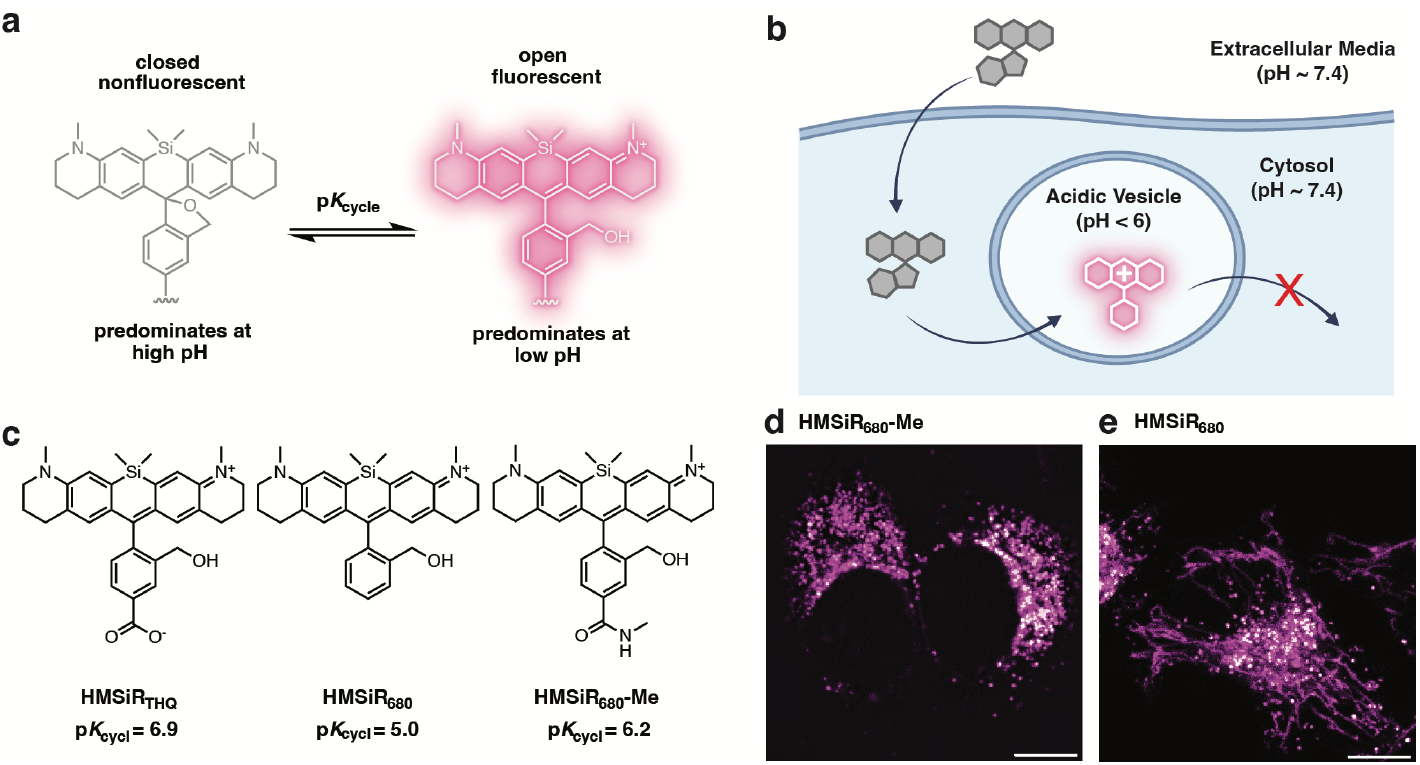
Design and initial characterization of HMSiR_680_-Me, a new silicon-rhodamine fluorophore that lights up at low pH to selectively image acidic organelles. (a) The emblematic pH-dependent on/off equilibrium of silicon rhodamine dyes that carry a hydroxymethyl group adjacent to the aromatic core (HMSiRs). (b) Strategy employed to target a pH-dependent HMSiR dye to acidic organelles. Although the dye may distribute throughout a cell, it will fluoresce only in regions where the pH is lower than pK_cycl_. (c) Structures and pK_cycl_ values of HMSiR_THQ_, HMSiR_680_, and HMSiR_680_-Me. (d-e) HeLa cells labeled with (d) HMSiR_680_-Me or (e) HMSiR_680_ and visualized using confocal microscopy. Scale bars = 10 μm.

Although HMSiR_THQ_ is too dim at neutral pH to support SRM, we reasoned that its unique photophysical features could translate into a versatile dye for imaging acidic organelles. We envisioned that upon protonation, HMSiR_THQ_ would become positively charged and membrane impermeable and thereby accumulate in acidic cellular compartments (Figure 1b). As reported, HMSiR_THQ_ contains a free carboxylate whose exceptionally low p*K*_a_ could confound lysosomal targeting. We designed and synthesized two HMSiR_THQ_ derivatives: HMSiR_680_, in which the carboxylate in HMSiR_THQ_ is deleted, and HMSiR_680_-Me, in which the carboxylate is replaced with a methyl amide (Figure 1c). Both derivatives demonstrated far-red fluorescence, with excitation maxima of 677 nm (HMSiR_680_) and 680 nm (HMSiR_680_-Me) (Figure S1a-b). HMSiR_680_-Me displayed a p*K*_cycl_ of 6.2 and when added to cells was localized to discrete punctae, as expected for organelles associated with the endocytic pathway (Figure 1d). In contrast, the p*K*_cycl_ of HMSiR_680_ was unexpectedly low (5.0) (Figure S1c) and when added to cells showed non-specific localization evidenced by elongated structures that matched signal localization of a mitochondrial GFP marker (Figures 1e and S1b). We concluded that HMSiR_680_-Me was a promising candidate for lysosome targeting and employed it exclusively in all subsequent experiments.

To confirm that the punctae observed in HMSiR_680_-Me-treated cells were acidic organelles, we performed colocalization experiments with *bona fide* markers for lysosomes (LAMP1-GFP), late endosomes (GFP-Rab7), early endosomes (GFP-Rab5), mitochondria (PDHA1-GFP), and the endoplasmic reticulum (GFP-KDEL). If the punctae observed in HMSiR_680_-Me-treated cells were acidic vesicles, we would expect strong colocalization with markers for lysosomes (pH ∼4.5) and late endosomes (pH ∼5.5), as the luminal pH of these organelles fall below the p*K*_cycl_ of HMSiR_680_-Me. In the same way, we would expect lower colocalization with markers for early endosomes (pH ∼6.5), mitochondria, and the endoplasmic reticulum, as the pH values of these organelles fall above the p*K*_cycl_ of HMSiR_680_-Me. HeLa cells expressing a single GFP-tagged organelle marker were treated with 500 nM HMSiR_680_-Me for 30 min, imaged using confocal microscopy, and the extent of colocalization evaluated by calculating Pearson’s correlation coefficients (PCC).^37^ Examination of the PCC values representing the colocalization of each GFP signal with that of HMSiR_680_-Me revealed strong colocalization with Lamp1 (0.75 ± 0.08) and Rab7 (0.80 ± 0.08), representing lysosomes and late endosomes, respectively. Significantly lower colocalization values were seen with Rab5 (0.47 ± 0.07), representing early endosomes. Only weak colocalization was detected with other organelle markers, indicating little or no localization within mitochondria or the endoplasmic reticulum (Figures 2a-b and S2a).

**Figure 2.**
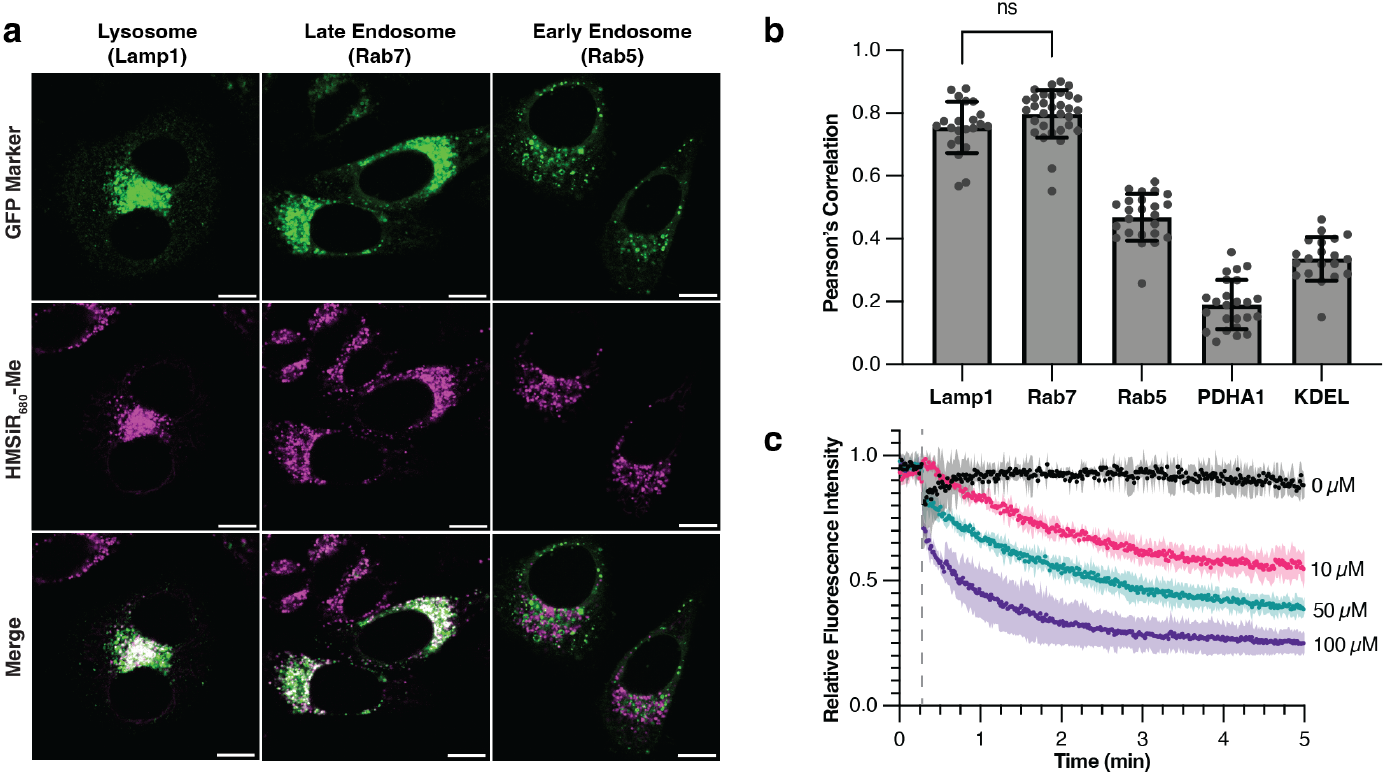
HMSiR_680_-Me lights up within acidic vesicles in live HeLa cells. (a) Representative confocal images of HeLa cells indicating the extent of colocalization between the signals due to HMSiR_680_-Me (magenta) and GFP organelle markers for lysosomes and early and late endosomes (green). Regions of the cell in which the magenta and green signals overlap appear white. Scale bars = 10 μm. (b) Plot illustrating the calculated Pearson’s correlation coefficients for n = 12 image sets for each organelle shown in part (a), plus those representing the extent of colocalization with signals due to mitochondria (PDHA1) and the ER (KDEL). “ns” indicates *p* > 0.05 (unpaired *t* test with Welch’s correction). (c) Fluorescence intensity of HeLa cells labeled with HMSiR_680_-Me and incubated with the indicated concentration of chloroquine diphosphate (CQ) for 5 min. The CQ solution was introduced to the cells 20 s into the 5 min imaging period (represented by dashed line). Data plotted in panel (c) represents 3 biological replicates; shaded area represents standard deviation.

Having confirmed that fluorescence due to HMSiR_680_-Me appeared selectively in late endosomes and lysosomes, organelles characterized by low luminal pH, we next sought to establish that the location-induced fluorescence was pH dependent. The luminal pH of late endosomes and lysosomes can be increased by treating cells with chloroquine diphosphate (CQ), a lysosomotropic compound that accumulates in acidic vesicles.^38,39^ We reasoned that if the fluorescence from HMSiR_680_-Me in lysosomes and late endosomes was due to the low luminal pH of these compartments, then the fluorescence signal associated with these organelles should decrease in the presence of CQ. Indeed, upon replacing imaging media with a CQ-containing solution, HeLa cells labeled with HMSiR_680_-Me showed a rapid and chloroquine concentration-dependent decrease in the fluorescence signal due to HMSiR_680_-Me (Figure 2c, Movie S1). Cells labeled with a Lamp1-GFP marker, whose fluorescence is not expected to be pH dependent, showed no decrease in signal in the presence of chloroquine (Figure S2b, Movie S1). Therefore, we concluded that the cellular fluorescence from HMSiR680-Me is indeed dependent on pH.

We next sought to investigate the utility of HMSiR680-Me for multicolor imaging. We were especially eager to evaluate whether HMSiR680-Me would support two-color imaging experiments that require only far-red excitation, which would be more benign to living cells. Two commonly used and commercially available far-red dyes are SiR and Cy5, which both have excitation maxima near 650 nm. Fluorophores with spectral properties similar to HMSiR_680_-Me, such as SiR_700_ and Yale_676sb_, are sufficiently spectrally separated from SiR to enable two-color experiments.^36,40^ Thus we envisioned that HMSiR_680_-Me could be used alongside SiR and Cy5 to enable two-color imaging using a simple coincubation protocol. To test this idea, we labeled HeLa cells with HMSiR_680_-Me and either Mitotracker Deep Red (MTDR), a Cy5-based dye, or SiR-DNA.^41^ In both cases we detected minimal bleedthrough of the HMSiR_680_-Me emission into the channel used to detect either MTDR or SiR; we calculated crosstalk values of 9% and 16% for MTDR and SiR-DNA, respectively. The bleed-through of the emission due to MTDR or SiR into the channel used to detect HMSiR_680_-Me was greater: 70% for MTDR and 25% for SiR-DNA (Figure S3). These results indicate that while spectral separation of both dyes with HMSiR_680_-Me is possible, the narrower emission spectrum of SiR makes it a better far-red two-color partner than Cy5. Performing linear unmixing^42-44^ gave spectral separation in both cases, and we ultimately achieved clear differentiation of lysosomes from either mitochondria (Figure 3a) or the nucleus (Figure 3b) in HeLa cells. Linear unmixing also allowed us to perform simultaneous three-color imaging of the nucleus, lysosomes, and mitochondria using SiR-DNA and HMSiR_680_-Me alongside commercially available MitoTracker Orange (Figures 3c and S4a) and PK Mito Orange (PKMO)^45^ (Figures 3d and S4b, Movie S2). These images clearly demonstrate the applicability of HMSiR_680_-Me for multicolor imaging by leveraging the nontoxic and infrequently used region of the visible spectrum above 650 nm.

**Figure 3.**
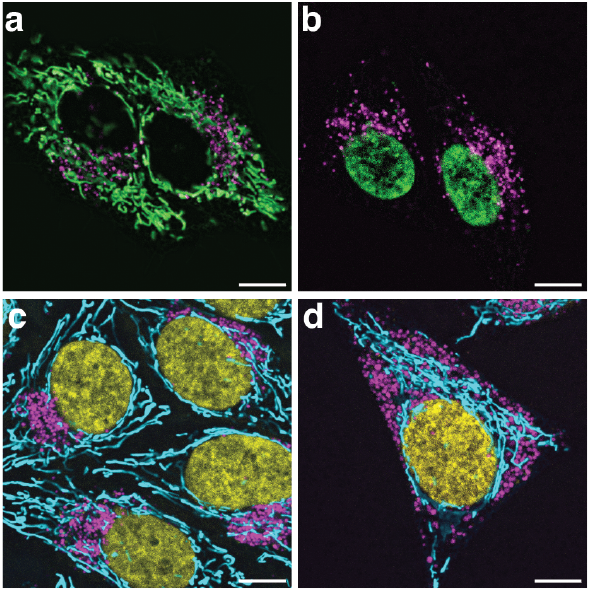
Two- and three-color confocal images of HeLa cells labeled with HMSiR_680_-Me and commercially available red and far-red dyes. HeLa cells labeled with (a) MitoTracker Deep Red (green) and HMSiR_680_-Me (magenta); (b) SiR-DNA (green) and HMSiR_680_-Me (magenta); (c) MitoTracker Orange (cyan), SiR-DNA (yellow), and HMSiR_680_-Me (magenta); (d) PKMito Orange (cyan), SiR-DNA (yellow), and HMSiR_680_-Me (magenta). Dyes were linearly unmixed using the Leica Stellaris Dye Separation tool. Scale bars = 10 μm. Dye concentrations were as follows: 500 nM HMSiR_680_-Me, 50 nM MitoTracker Deep Red, 1 μM SiR-DNA, 100 nM MitoTracker Orange, 1x PKMito Orange.

As mentioned previously, an advantage of silicon rhodamine fluorophores is their ability to withstand the high-intensity depletion laser required for stimulated emission depletion (STED) microscopy. To investigate whether HMSiR_680_-Me would support STED microscopy, and to visually confirm that its fluorescence was localized to the lysosomal lumen, we labeled HeLa cells with Lamp1-SiR, using LAMP1-HaloTag and SiR-CA, as well as HMSiR_680_-Me.^46^

Because the HaloTag is appended to the cytosolic C-terminus of Lamp1, we expect to see clear differentiation of the luminal signals from HMSiR_680_-Me and the membrane signals from SiR. Indeed, when imaged via STED, the SiR signal (green) appears as a border around the HMSiR_680_-Me signal (magenta) in many organelles, confirmed quantitatively with a line profile (Figure 4). These images further support the hypothesis that the fluorescence due to HMSiR_680_-Me localizes within the acidic lumen of endocytic vesicles. We also confirmed that this level of resolution, particularly in the Lamp1-SiR channel, could not be achieved with confocal microscopy, further enforcing the need for improved dyes for STED (Figure S5).

**Figure 4.**
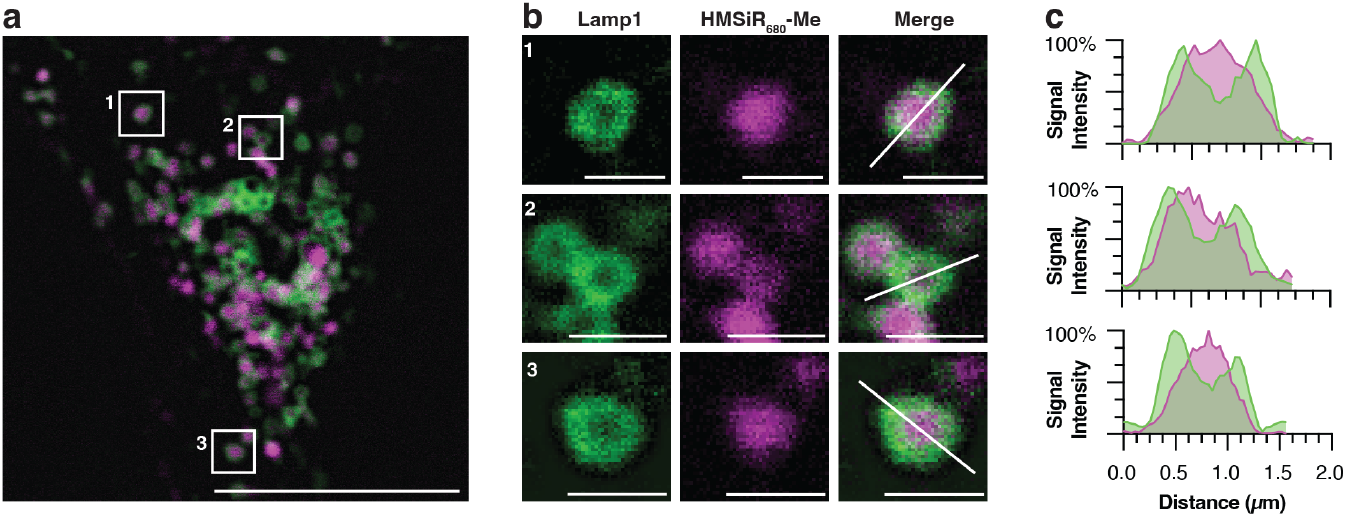
STED demonstrates that HMSiR_680_-Me localizes to the lysosomal lumen in HeLa cells. (a) Cells labeled with HMSiR_680_-Me (magenta) and LAMP1-HaloTag and SiR-chloroalkane (green) and imaged via STED microscopy. Scale bar = 10 μm. (b) Insets from panel (a). Scale bars = 1 μm. (c) Line profile diagrams along the lines shown in panel (b). Maximum intensity for each channel was scaled. Dye concentrations were 500 nM for HMSiR_680_-Me and 2 μM for SiR-chloroalkane.

Another imaging challenge presented by lysosome function is their rapid movement. Lysosomes move quickly (1 μm/s) along microtubules, and their motility is an established metric for cell health.^8^ Measuring lysosomal dynamics is difficult because it requires imaging many frames at a high frame rate. Thus, the dye employed must be both sufficiently bright to support a high frame rate as well as sufficiently photostable to support data acquisition over many frames.^47^ To assess whether HMSiR_680_-Me would support dynamic lysosomal imaging, we compared it to LTDR, a commonly used and commercially available far-red lysosome probe. When evaluated at an equivalent concentration (500 nM) in vitro (0.2 M phosphate buffer, pH = 4.5), HMSiR_680_-Me was 25-fold more photostable than LTDR, retaining 54% of its initial intensity after 2 h compared to 2.5% for LTDR (Figure 5a). The loss of fluorescence due to LTDR followed a one-phase decay, whereas the loss of fluorescence due to HMSiR_680_-Me followed a two-phase decay. When HeLa cells were labeled with HMSiR_680_-Me or LTDR at 500 nM and imaged with a frame rate of 1 fps, the dyes bleached at a similar rate, with 49% or 53% of the initial signal remaining after 1 h for HMSiR_680_-Me and LTDR, respectively. Notably, to avoid overexposure, cells labeled with LTDR were imaged with only 5% laser power compared to 20% for HMSiR_680_-Me, yet LTDR lost nearly the same amount of signal as HMSiR_680_-Me. To compare the extent of photobleaching under equivalent conditions, we decreased the concentration of LTDR to 50 nM and increased the laser power to 20%, and under these conditions we observed only 16% of the initial LTDR signal remaining after 1h, compared to 49% for HMSiR_680_-Me (Figure 5b). Taken together, these results suggest that HMSiR_680_-Me is more photostable than LTDR in cells as well as in vitro.

**Figure 5.**
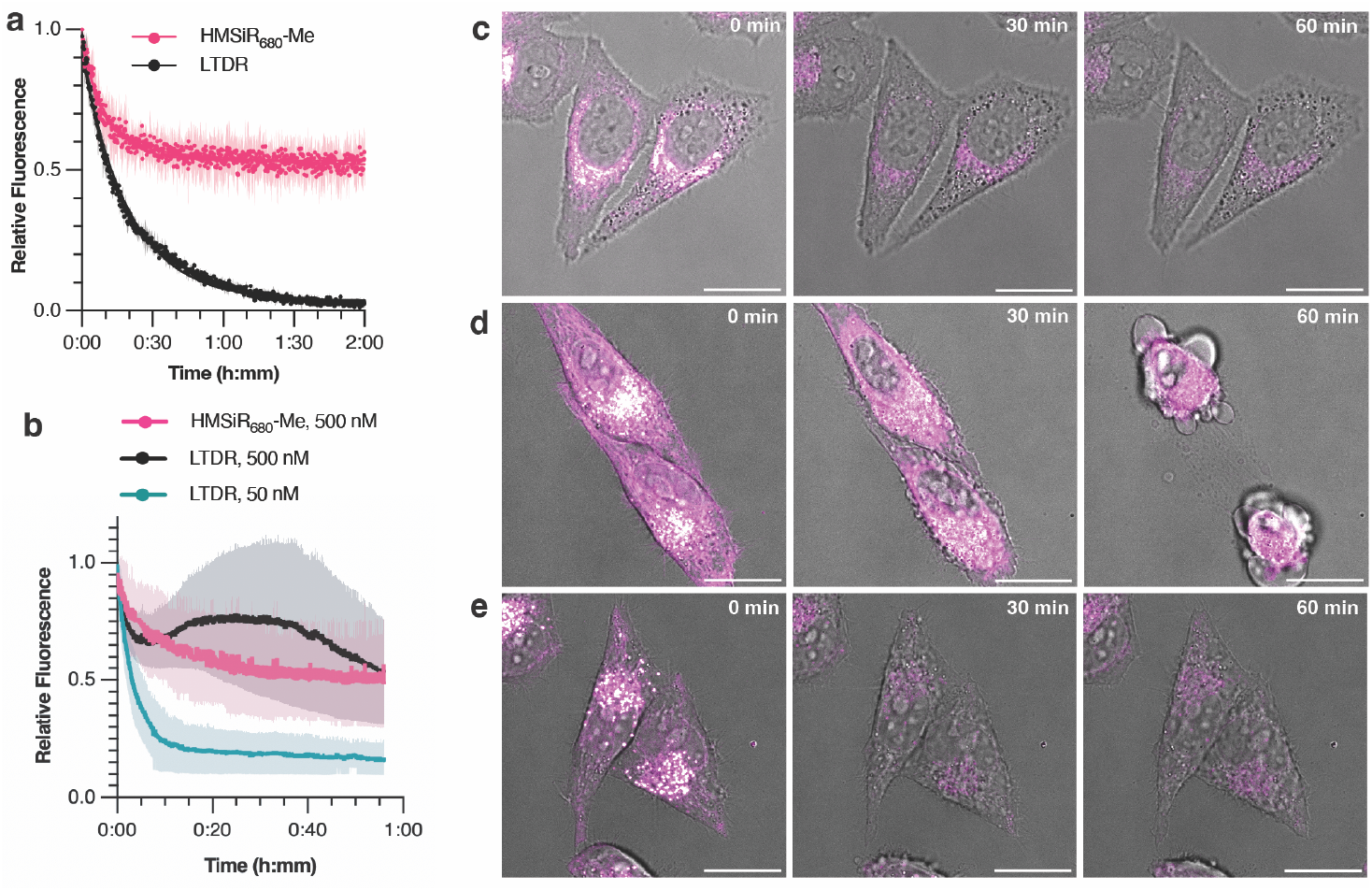
HMSiR_680_-Me shows improved photostability and decreased cytotoxicity compared to LTDR. (a) Fluorescence intensity over time for 500 nM HMSiR_680_-Me and LTDR in 0.2 M phosphate buffer, pH = 4.5. Data represents 3 technical replicates. (b) Fluorescence intensity over time for HeLa cells labeled with (c) 500 nM HMSiR_680_-Me, (d) 500 nM LTDR, and (e) 50 nM LTDR. Data plotted in panel (b) represents 3 biological replicates; shaded area represents standard deviation. Scale bars = 20 μm.

HMSiR_680_-Me also appeared to be more benign to cell health when imaging for many frames, even when used at a relatively high concentration. Cells labeled with 500 nM LTDR showed significant off-target labeling and blebbing during the course of the experiment, indicating a toxic side-effect. In contrast, cells labeled with 500 nM HMSiR_680_-Me and 50 nM LTDR remained healthy throughout the time lapse (Figure 5c-e, Movie S3). Upon close inspection, we also observed a decrease in lysosome motility over the course of the experiment when cells were labeled with LTDR. To quantify the effect of both dyes on lysosome motility, we imaged cells labeled with 500 nM HMSiR_680_-Me or 50 nM LTDR for 1000 frames at 1 fps and 20% laser power (Movie S4). For three biological replicates of each dye, we plotted lysosome speed over time and performed a linear regression, and we found the average slope to be significantly nonzero only for LTDR (Figure S6a-b). To visualize the data another way, we compared the speed of lysosomes detected within the first 50 frames with those detected during the last 50 frames, and we observed a significant decrease in speed only when cells were labeled with LTDR (Figure S6c). These data indicate that HMSiR_680_-Me enables confocal imaging of lysosomes over many frames at a high frame rate without risk of deleterious effects on organelle motility and cell health.

We then showed that HMSiR_680_-Me is useful for other time lapse imaging applications. To demonstrate its applicability in long imaging experiments, we decreased the frame rate to 1 frame every 2 minutes and imaged HMSiR_680_-Melabeled HeLa cells for 16 hours with excellent signal retention and continued cell division (Figure S7, Movie S5). Using a 775 nm depletion laser, we were also able to achieve STED imaging of lysosomes labeled with HMSiR_680_-Me at an exceptionally fast frame rate of 1.5 fps for at least 1000 frames with a pixel size of only 50 nm (Figure S8, Movie S6). Together, these time lapse images suggest that HMSiR_680_-Me can be applied to a broad range of biological experiments, whether the system of interest is multicellular or subcellular.

Finally, the ability to image lysosomes alongside other organelles at super resolution could enable the study of organelle-organelle interactions below the diffraction limit. As mentioned, lysosomes aid in the transfer of nutrients and metabolites to other organelles, contributing to normal cell function as well as cellular dysfunction in disease. Specifically, defective mitochondria–lysosome interactions are associated with neurodegenerative disease and cancer.^48-50^ Therefore, we aimed to image this interaction using STED, which would require two dyes each targeting one of the two organelles. We previously confirmed that the fluorescence signal from HMSiR_680_-Me and SiR could be effectively separated with linear unmixing, and we anticipated that their excitation spectra would overlap sufficiently to facilitate simultaneous excitation with a single laser. Thus, to enable multiplexed imaging of lysosomes and mitochondria, we treated HeLa cells with both HMSiR_680_-Me as well as recently reported MAO-SiR, an inner mitochondrial membrane (IMM) High Density Environmentally Sensitive (HIDE) probe, which results from in cellulo reaction of MAO-N_3_ with SiR-DBCO.^51^ MAO-SiR images the IMM selectively, continuously, and at super resolution for extended periods of time without extensive photobleaching or toxicity. Treated cells excited with a single 645 nm laser and signal were monitored in multiple detection windows between 655 and 770 nm. After linear unmixing, we observed distinct labeling of elongated mitochondrial structures and acidic vesicles that lasted for up to 300 confocal imaging frames (Figure 6a-b, Movie S7). We applied the same approach to obtain images of detailed IMM structure alongside the lysosome for up to 60 STED imaging frames (Figure 6c-d, Movie S8). These two-color images were collected with only a single excitation laser and single depletion laser per frame, significantly improving temporal resolution and photostability compared to a two-color image where dyes are excited and depleted sequentially. At 0.385 fps and a pixel size of only 30 nm, this multicolor time lapse is an excellent demonstration of how HMSiR_680_-Me can be multiplexed with other far-red dyes to image organelle-organelle interactions well below the diffraction limit without sacrificing temporal resolution.

**Figure 6.**
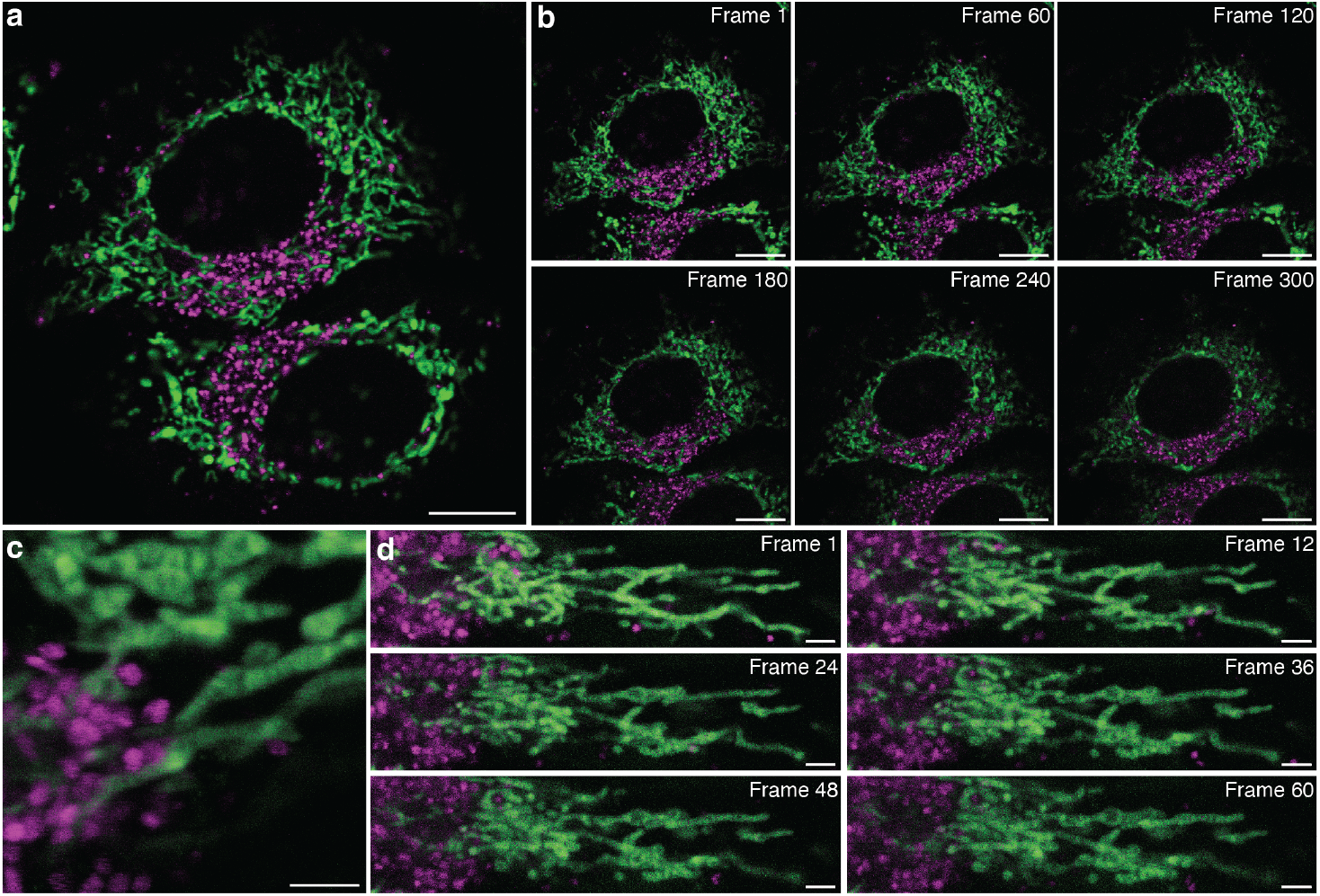
HMSiR_680_-Me (magenta) and MAO-N_3_/SiR-DBCO (green) can label the lysosome and inner mitochondrial membrane of HeLa cells in a multiplexed imaging experiment. (a) Confocal, scale bar = 5 μm. (b) Confocal time lapse, scale bars = 5 μm. (c) STED, scale bar = 2 μm. (d) STED time lapse, scale bars = 2 μm. Dyes were linearly unmixed using the Leica Stellaris Dye Separation tool. Dye concentrations were 1 μM for HMSiR_680_-Me and 100 nM for SiR-DBCO.

## CONCLUSIONS

HMSiR_680_-Me is a superior small-molecule fluorophore for lysosomal imaging that specifically labels acidic organelles in a pH-dependent manner. It demonstrates excellent photostability and no detrimental effects on cell health and organelle motility. Here we showcase its utility in a wide range of live-cell experiments, including continual overnight imaging and time lapse STED imaging with a frame rate faster than 1 fps. Its 680 nm excitation wavelength is spectrally separate from other commonly used far-red dyes such as SiR and Cy5, enabling two- and three-color imaging using only far-red excitation. Furthermore, HMSiR_680_-Me is compatible with multicolor STED, and when used alongside SiR, enables two-color imaging of lysosomal dynamics beside detailed IMM structure. Importantly, these time lapse STED images were collected with only a single excitation and depletion exposure per frame, significantly improving the temporal resolution and photostability compared to sequential two-color imaging. As previously mentioned, silicon rhodamine fluorophores are highly modular, and the novel targeting strategy employed to design HMSiR_680_-Me could be used to develop hydroxymethyl rhodamine-based acidic organelle probes with a wide range of spectral and chemical properties. Furthermore, we envision that the methyl amide on the pendant ring of HMSiR_680_-Me could provide a convenient handle for conjugation to biomolecules of interest. We expect that HMSiR_680_-Me will prove to be an indispensable tool for studying the role of lysosomes in health and disease.

## ASSOCIATED CONTENT

### Supporting Information

The Supporting Information is available free of charge on the ACS Publications website.

Supplementary figures, movie descriptions, experimental methods, and characterization of new compounds (PDF)

Movie S1: HMSiR_680_-Me is pH-dependent in live cells (MOV)

Movie S2: HMSiR_680_-Me can be used with other red and far-red dyes in multicolor imaging experiments (AVI)

Movie S3: HMSiR_680_-Me photobleaches slowly and does not affect cell health (MOV)

Movie S4: HMSiR_680_-Me does not affect lysosomal motility (MOV)

Movie S5: HMSiR_680_-Me is compatible with overnight confocal imaging (MOV)

Movie S6: HMSiR_680_-Me is compatible with time lapse STED imaging with excellent temporal resolution (AVI)

Movie S7: HMSiR_680_-Me can be used with SiR in a two-color multiplexed imaging experiment (AVI)

Movie S8: HMSiR_680_-Me can be used with SiR in a two-color multiplexed STED imaging experiment (AVI)

## AUTHOR INFORMATION

## ACKNOWLEDGMENT

We are grateful to all members of the Schepartz lab for support and helpful discussions. Research reported in this publication was supported by the National Institute of General Medical Sciences of the National Institutes of Health under Award Number R35GM134963. L.L. was supported in part by a National Science Foundation Graduate Research Fellowship (Grant No. 2146752). Any opinions, findings, and conclusions or recommendations expressed in this material are those of the authors and do not necessarily reflect the views of the National Science Foundation. Fluorescence spectra and absolute quantum yield measurements were performed at the Molecular Foundry (LBNL), which is supported by the Office of Science, Office of Basic Energy Sciences, of the U.S. Department of Energy under Contract No. DE-AC02-05CH11231. We thank Dr. Hasan Celik and UC Berkeley’s NMR facility in the College of Chemistry (CoC-NMR) for spectroscopic assistance. Instruments in the CoC-NMR are supported in part by NIH S10OD024998. The plasmid used to express CMV-LAMP1-HaloTag plasmid was a gift from Jennifer Lippincott-Schwartz (HHMI-Janelia).

## REFERENCES

(1) Lawrence, R. E.; Zoncu, R. The Lysosome as a Cellular Centre for Signalling, Metabolism and Quality Control. Nat. Cell Biol. 2019, 21 (2), 133–142. https://doi.org/10.1038/s41556-018-0244-7.

(2) Banik, S. M.; Pedram, K.; Wisnovsky, S.; Ahn, G.; Riley, N. M.; Bertozzi, C. R. Lysosome-Targeting Chimaeras for Degradation of Extracellular Proteins. Nature 2020, 584 (7820), 291–297. https://doi.org/10.1038/s41586-020-2545-9.

(3) Pance, K.; Gramespacher, J. A.; Byrnes, J. R.; Salangsang, F.; Serrano, J.-A. C.; Cotton, A. D.; Steri, V.; Wells, J. A. Modular Cytokine Receptor-Targeting Chimeras for Targeted Degradation of Cell Surface and Extracellular Proteins. Nat. Biotechnol. 2023, 41 (2), 273–281. https://doi.org/10.1038/s41587-022-01456-2.

(4) Liu, G. Y.; Sabatini, D. M. MTOR at the Nexus of Nutrition, Growth, Ageing and Disease. Nat. Rev. Mol. Cell Biol. 2020, 21 (4), 183–203. https://doi.org/10.1038/s41580-019-0199-y.

(5) Ballabio, A.; Bonifacino, J. S. Lysosomes as Dynamic Regulators of Cell and Organismal Homeostasis. Nat. Rev. Mol. Cell Biol. 2020, 21 (2), 101–118. https://doi.org/10.1038/s41580-019-0185-4.

(6) Medoh, U. N.; Chen, J. Y.; Abu-Remaileh, M. Lessons from Metabolic Perturbations in Lysosomal Storage Disorders for Neurodegeneration. Curr. Opin. Syst. Biol. 2022, 29, 100408. https://doi.org/10.1016/j.coisb.2021.100408.

(7) Cranfill, P. J.; Sell, B. R.; Baird, M. A.; Allen, J. R.; Lavagnino, Z.; de Gruiter, H. M.; Kremers, G.-J.; Davidson, M. W.; Ustione, A.; Piston, D. W. Quantitative Assessment of Fluorescent Proteins. Nat. Methods 2016, 13 (7), 557–562. https://doi.org/10.1038/nmeth.3891.

(8) Barral, D. C.; Staiano, L.; Guimas Almeida, C.; Cutler, D. F.; Eden, E. R.; Futter, C. E.; Galione, A.; Marques, A. R. A.; Medina, D. L.; Napolitano, G.; Settembre, C.; Vieira, O. V.; Aerts, J. M. F. G.; Atakpa-Adaji, P.; Bruno, G.; Capuozzo, A.; De Leonibus, E.; Di Malta, C.; Escrevente, C.; Esposito, A.; Grumati, P.; Hall, M. J.; Teodoro, R. O.; Lopes, S. S.; Luzio, J. P.; Monfregola, J.; Montefusco, S.; Platt, F. M.; Polishchuck, R.; De Risi, M.; Sambri, I.; Soldati, C.; Seabra, M. C. Current Methods to Analyze Lysosome Morphology, Positioning, Motility and Function. Traffic 2022, 23 (5), 238–269. https://doi.org/10.1111/tra.12839.

(9) de Araujo, M. E. G.; Liebscher, G.; Hess, M. W.; Huber, L. A. Lysosomal Size Matters. Traffic 2020, 21 (1), 60–75. https://doi.org/10.1111/tra.12714.

(10) Dadina, N.; Tyson, J.; Zheng, S.; Lesiak, L.; Schepartz, A. Imaging Organelle Membranes in Live Cells at the Nanoscale with Lipid-Based Fluorescent Probes. Curr. Opin. Chem. Biol. 2021, 65, 154–162. https://doi.org/10.1016/j.cbpa.2021.09.003.

(11) Grimm, J. B.; Lavis, L. D. Caveat Fluorophore: An Insiders’ Guide to Small-Molecule Fluorescent Labels. Nat. Methods 2022, 19 (2), 149–158. https://doi.org/10.1038/s41592-021-01338-6.

(12) Chu, L.; Tyson, J.; Shaw, J. E.; Rivera-Molina, F.; Koleske, A. J.; Schepartz, A.; Toomre, D. K. Two-Color Nanoscopy of Organelles for Extended Times with HIDE Probes. Nat. Commun. 2020, 11 (1), 4271. https://doi.org/10.1038/s41467-020-17859-1.

(13) Chiu, D.-C.; Baskin, J. M. Imaging and Editing the Phospholipidome. Acc. Chem. Res. 2022, 55 (21), 3088–3098. https://doi.org/10.1021/acs.accounts.2c00510.

(14) Klymchenko, A. S. Fluorescent Probes for Lipid Membranes: From the Cell Surface to Organelles. Acc. Chem. Res. 2023, 56 (1), 1–12. https://doi.org/10.1021/acs.accounts.2c00586.

(15) Los, G. V.; Encell, L. P.; McDougall, M. G.; Hartzell, D. D.; Karassina, N.; Zimprich, C.; Wood, M. G.; Learish, R.; Ohana, R. F.; Urh, M.; Simpson, D.; Mendez, J.; Zimmerman, K.; Otto, P.; Vidugiris, G.; Zhu, J.; Darzins, A.; Klaubert, D. H.; Bulleit, R. F.; Wood, K. V. HaloTag: A Novel Protein Labeling Technology for Cell Imaging and Protein Analysis. ACS Chem. Biol. 2008, 3 (6), 373–382. https://doi.org/10.1021/cb800025k.

(16) Wang, L.; Frei, M. S.; Salim, A.; Johnsson, K. Small-Molecule Fluorescent Probes for Live-Cell Super-Resolution Microscopy. J. Am. Chem. Soc. 2019, 141 (7), 2770–2781. https://doi.org/10.1021/jacs.8b11134.

(17) Erdmann, R. S.; Baguley, S. W.; Richens, J. H.; Wissner, R. F.; Xi, Z.; Allgeyer, E. S.; Zhong, S.; Thompson, A. D.; Lowe, N.; Butler, R.; Bewersdorf, J.; Rothman, J. E.; St Johnston, D.; Schepartz, A.; Toomre, D. Labeling Strategies Matter for Super-Resolution Microscopy: A Comparison between HaloTags and SNAP-Tags. Cell Chem. Biol. 2019, 26 (4), 584-592.e6. https://doi.org/10.1016/j.chembiol.2019.01.003.

(18) Richardson, D. S.; Gregor, C.; Winter, F. R.; Urban, N. T.; Sahl, S. J.; Willig, K. I.; Hell, S. W. SRpHi Ratiometric PH Biosensors for Super-Resolution Microscopy. Nat. Commun. 2017, 8 (1), 577. https://doi.org/10.1038/s41467-017-00606-4.

(19) He, H.; Ye, Z.; Zheng, Y.; Xu, X.; Guo, C.; Xiao, Y.; Yang, W.; Qian, X.; Yang, Y. Super-Resolution Imaging of Lysosomes with a Nitroso-Caged Rhodamine. Chem. Commun. 2018, 54 (23), 2842–2845. https://doi.org/10.1039/C7CC08886H.

(20) Xue, Z.; Wang, S.; Li, J.; Chen, X.; Han, J.; Han, S. Bifunctional Super-Resolution Imaging Probe with Acidity-Independent Lysosome-Retention Mechanism. Anal. Chem. 2018, 90 (19), 11393–11400. https://doi.org/10.1021/acs.analchem.8b02365.

(21) Wang, C.; Taki, M.; Kajiwara, K.; Wang, J.; Yamaguchi, S. Phosphole-Oxide-Based Fluorescent Probe for Super-Resolution Stimulated Emission Depletion Live Imaging of the Lysosome Membrane. ACS Mater. Lett. 2020, 2 (7), 705–711. https://doi.org/10.1021/acsmaterialslett.0c00147.

(22) Ye, Z.; Yang, W.; Wang, C.; Zheng, Y.; Chi, W.; Liu, X.; Huang, Z.; Li, X.; Xiao, Y. Quaternary Piperazine-Substituted Rhodamines with Enhanced Brightness for Super-Resolution Imaging. J. Am. Chem. Soc. 2019, 141 (37), 14491–14495. https://doi.org/10.1021/jacs.9b04893.

(23) Yadav, A.; Rao, C.; Nandi, C. K. Fluorescent Probes for Super-Resolution Microscopy of Lysosomes. ACS Omega 2020, 5 (42), 26967–26977. https://doi.org/10.1021/acsomega.0c04018.

(24) Chen, Q.; Fang, H.; Shao, X.; Tian, Z.; Geng, S.; Zhang, Y.; Fan, H.; Xiang, P.; Zhang, J.; Tian, X.; Zhang, K.; He, W.; Guo, Z.; Diao, J. A Dual-Labeling Probe to Track Functional Mitochondria– Lysosome Interactions in Live Cells. Nat. Commun. 2020, 11 (1), 6290. https://doi.org/10.1038/s41467-020-20067-6.

(25) Ye, Z.; Zheng, Y.; Peng, X.; Xiao, Y. Surpassing the Background Barrier for Multidimensional Single-Molecule Localization Super-Resolution Imaging: A Case of Lysosome-Exclusively Turn-on Probe. Anal. Chem. 2022, 94 (22), 7990–7995. https://doi.org/10.1021/acs.analchem.2c00987.

(26) Qiao, Q.; Liu, W.; Chen, J.; Wu, X.; Deng, F.; Fang, X.; Xu, N.; Zhou, W.; Wu, S.; Yin, W.; Liu, X.; Xu, Z. An Acid-Regulated Self-Blinking Fluorescent Probe for Resolving Whole-Cell Lysosomes with Long-Term Nanoscopy. Angew. Chem. 2022, 134 (21), e202202961. https://doi.org/10.1002/ange.202202961.

(27) Wu, L.; Li, X.; Ling, Y.; Huang, C.; Jia, N. Morpholine Derivative-Functionalized Carbon Dots-Based Fluorescent Probe for Highly Selective Lysosomal Imaging in Living Cells. ACS Appl. Mater. Interfaces 2017, 9 (34), 28222–28232. https://doi.org/10.1021/acsami.7b08148.

(28) Zhang, Y.; Song, K.-H.; Tang, S.; Ravelo, L.; Cusido, J.; Sun, C.; Zhang, H. F.; Raymo, F. M. Far-Red Photoactivatable BODIPYs for the Super-Resolution Imaging of Live Cells. J. Am. Chem. Soc. 2018, 140 (40), 12741–12745. https://doi.org/10.1021/jacs.8b09099.

(29) Fan, M.; An, H.; Wang, C.; Huo, S.; Wang, T.; Cui, X.; Zhang, D. STED Imaging the Dynamics of Lysosomes by Dually Fluorogenic Si-Rhodamine. Chem. – Eur. J. 2021, 27 (37), 9620–9626. https://doi.org/10.1002/chem.202100623.

(30) Ogasawara, H.; Tanaka, Y.; Taki, M.; Yamaguchi, S. Late-Stage Functionalisation of Alkyne-Modified Phospha-Xanthene Dyes: Lysosomal Imaging Using an off–on–off Type of PH Probe. Chem. Sci. 2021, 12 (22), 7902–7907. https://doi.org/10.1039/D1SC01705E.

(31) Grimm, J. B.; English, B. P.; Chen, J.; Slaughter, J. P.; Zhang, Z.; Revyakin, A.; Patel, R.; Macklin, J. J.; Normanno, D.; Singer, R. H.; Lionnet, T.; Lavis, L. D. A General Method to Improve Fluorophores for Live-Cell and Single-Molecule Microscopy. Nat. Methods 2015, 12 (3), 244–250. https://doi.org/10.1038/nmeth.3256.

(32) Grimm, J. B.; Muthusamy, A. K.; Liang, Y.; Brown, T. A.; Lemon, W. C.; Patel, R.; Lu, R.; Macklin, J. J.; Keller, P. J.; Ji, N.; Lavis, L. D. A General Method to Fine-Tune Fluorophores for Live-Cell and in Vivo Imaging. Nat. Methods 2017, 14 (10), 987–994. https://doi.org/10.1038/nmeth.4403.

(33) Zheng, Q.; Ayala, A. X.; Chung, I.; Weigel, A. V.; Ranjan, A.; Falco, N.; Grimm, J. B.; Tkachuk, A. N.; Wu, C.; Lippincott-Schwartz, J.; Singer, R. H.; Lavis, L. D. Rational Design of Fluorogenic and Spontaneously Blinking Labels for Super-Resolution Imaging. ACS Cent. Sci. 2019, 5 (9), 1602–1613. https://doi.org/10.1021/acscentsci.9b00676.

(34) Koide, Y.; Urano, Y.; Hanaoka, K.; Terai, T.; Nagano, T. Evolution of Group 14 Rhodamines as Platforms for Near-Infrared Fluorescence Probes Utilizing Photoinduced Electron Transfer. ACS Chem. Biol. 2011, 6 (6), 600–608. https://doi.org/10.1021/cb1002416.

(35) Koide, Y.; Urano, Y.; Hanaoka, K.; Piao, W.; Kusakabe, M.; Saito, N.; Terai, T.; Okabe, T.; Nagano, T. Development of NIR Fluorescent Dyes Based on Si–Rhodamine for in Vivo Imaging. J. Am. Chem. Soc. 2012, 134 (11), 5029–5031. https://doi.org/10.1021/ja210375e.

(36) Tyson, J.; Hu, K.; Zheng, S.; Kidd, P.; Dadina, N.; Chu, L.; Toomre, D.; Bewersdorf, J.; Schepartz, A. Extremely Bright, Near-IR Emitting Spontaneously Blinking Fluorophores Enable Ratiometric Multicolor Nanoscopy in Live Cells. ACS Cent. Sci. 2021, 7 (8), 1419–1426. https://doi.org/10.1021/acscentsci.1c00670.

(37) Manders, E. M. M.; Verbeek, F. J.; Aten, J. A. Measurement of Co-Localization of Objects in Dual-Colour Confocal Images. J. Microsc. 1993, 169 (3), 375–382. https://doi.org/10.1111/j.1365-2818.1993.tb03313.x.

(38) Ohkuma, S.; Poole, B. Fluorescence Probe Measurement of the Intralysosomal PH in Living Cells and the Perturbation of PH by Various Agents. Proc. Natl. Acad. Sci. U. S. A. 1978, 75 (7), 3327–3331. https://doi.org/10.1073/pnas.75.7.3327.

(39) Halcrow, P. W.; Geiger, J. D.; Chen, X. Overcoming Chemoresistance: Altering PH of Cellular Compartments by Chloroquine and Hydroxychloroquine. Front. Cell Dev. Biol. 2021, 9, 627639. https://doi.org/10.3389/fcell.2021.627639.

(40) Lukinavičius, G.; Reymond, L.; Umezawa, K.; Sallin, O.; D’Este, E.; Göttfert, F.; Ta, H.; Hell, S. W.; Urano, Y.; Johnsson, K. Fluorogenic Probes for Multicolor Imaging in Living Cells. J. Am. Chem. Soc. 2016, 138 (30), 9365–9368. https://doi.org/10.1021/jacs.6b04782.

(41) Lukinavičius, G.; Blaukopf, C.; Pershagen, E.; Schena, A.; Reymond, L.; Derivery, E.; Gonzalez-Gaitan, M.; D’Este, E.; Hell, S. W.; Wolfram Gerlich, D.; Johnsson, K. SiR–Hoechst Is a Far-Red DNA Stain for Live-Cell Nanoscopy. Nat. Commun. 2015, 6 (1), 8497. https://doi.org/10.1038/ncomms9497.

(42) Zimmermann, T. Spectral Imaging and Linear Unmixing in Light Microscopy. In Microscopy Techniques: -/-; Rietdorf, J., Ed.; Advances in Biochemical Engineering; Springer: Berlin, Heidelberg, 2005; pp 245–265. https://doi.org/10.1007/b102216.

(43) Valm, A. M.; Cohen, S.; Legant, W. R.; Melunis, J.; Hershberg, U.; Wait, E.; Cohen, A. R.; Davidson, M. W.; Betzig, E.; Lippincott-Schwartz, J. Applying Systems-Level Spectral Imaging and Analysis to Reveal the Organelle Interactome. Nature 2017, 546 (7656), 162–167. https://doi.org/10.1038/nature22369.

(44) Acuña-Rodriguez, J. P.; Mena-Vega, J. P.; Argüello-Miranda, O. Live-Cell Fluorescence Spectral Imaging as a Data Science Challenge. Biophys. Rev. 2022, 14 (2), 579–597. https://doi.org/10.1007/s12551-022-00941-x.

(45) Liu, T.; Stephan, T.; Chen, P.; Keller-Findeisen, J.; Chen, J.; Riedel, D.; Yang, Z.; Jakobs, S.; Chen, Z. Multi-Color Live-Cell STED Nanoscopy of Mitochondria with a Gentle Inner Membrane Stain. Proc. Natl. Acad. Sci. 2022, 119 (52), e2215799119. https://doi.org/10.1073/pnas.2215799119.

(46) Liao, Y.-C.; Fernandopulle, M. S.; Wang, G.; Choi, H.; Hao, L.; Drerup, C. M.; Patel, R.; Qamar, S.; Nixon-Abell, J.; Shen, Y.; Meadows, W.; Vendruscolo, M.; Knowles, T. P. J.; Nelson, M.; Czekalska, M. A.; Musteikyte, G.; Gachechiladze, M. A.; Stephens, C. A.; Pasolli, H. A.; Forrest, L. R.; St George-Hyslop, P.; Lippincott-Schwartz, J.; Ward, M. E. RNA Granules Hitchhike on Lysosomes for Long-Distance Transport, Using Annexin A11 as a Molecular Tether. Cell 2019, 179 (1), 147-164.e20. https://doi.org/10.1016/j.cell.2019.08.050.

(47) Qiu, K.; Yadav, A.; Tian, Z.; Guo, Z.; Shi, D.; Nandi, C. K.; Diao, J. Ultralong-Term Super-Resolution Tracking of Lysosomes in Brain Organoids by Near-Infrared Noble Metal Nanoclusters. ACS Mater. Lett. 2022, 4 (9), 1565–1573. https://doi.org/10.1021/acsmaterialslett.2c00436.

(48) Wong, Y. C.; Ysselstein, D.; Krainc, D. Mitochondria– Lysosome Contacts Regulate Mitochondrial Fission via RAB7 GTP Hydrolysis. Nature 2018, https://doi.org/10.1038/nature25486. 554 (7692), p382–386.

(49) Wong, Y. C.; Kim, S.; Peng, W.; Krainc, D. Regulation and Function of Mitochondria–Lysosome Membrane Contact Sites in Cellular Homeostasis. Trends Cell Biol. 2019, 29 (6), 500–513. https://doi.org/10.1016/j.tcb.2019.02.004.

(50) Deus, C. M.; Yambire, K. F.; Oliveira, P. J.; Raimundo, N. Mitochondria–Lysosome Crosstalk: From Physiology to Neurodegeneration. Trends Mol. Med. 2020, 26 (1), 71–88. https://doi.org/10.1016/j.molmed.2019.10.009.

(51) Zheng, S.; Dadina, N.; Mozumdar, D.; Lesiak, L.; Martinez, K. N.; Miller, E.; Schepartz, A. Long-Term Super-Resolution Inner Mitochondrial Membrane Imaging with a Lipid Probe. Nat. Chem. Biol. 2023, In Press.

