## Supplementary Info for "A Bright, Photostable Dye that Enables Multicolor, Time Lapse, and Super-Resolution Imaging of Acidic Organelles"

Supporting Information

Supporting Information: Table of Contents

### I. Supplementary Figures

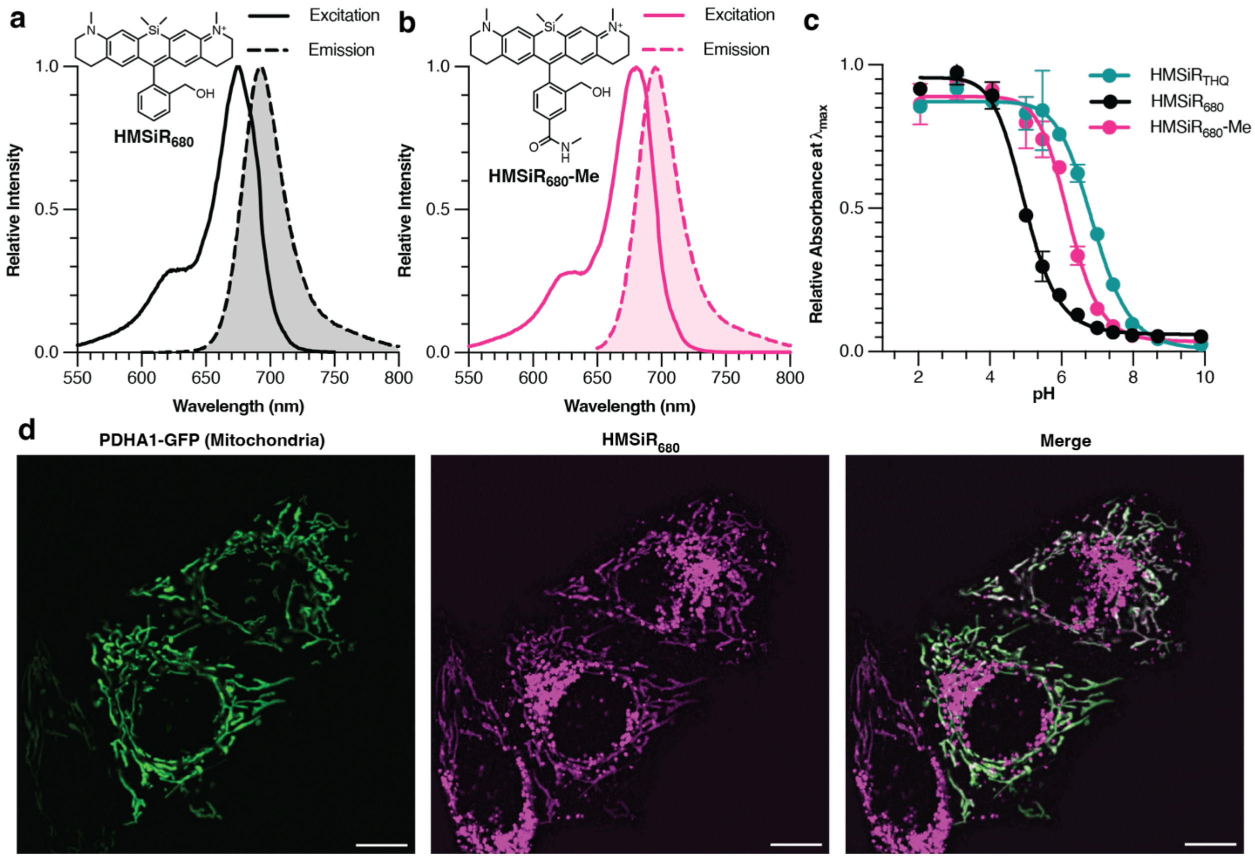

**Figure S1.** HMSiR$_{680}$-Me is a better far-red probe for acidic vesicles than HMSiR$_{680}$. Excitation and emission spectra for newly reported molecules (a) HMSiR$_{680}$ and (b) HMSiR$_{680}$-Me. (c) pH-dependent change in absorbance of 15 µM HMSiR$_{THQ}$ (monitored at 680 nm), HMSiR$_{680}$ (monitored at 677 nm), and HMSiR$_{680}$-Me (monitored at 680 nm) in 0.2 M phosphate buffer at room temperature. The absorbance intensity for each molecule was measured at its maximum absorbance wavelength in phosphate-buffered solution, pH = 4.5. (d) Confocal images of HeLa cells transfected with PDHA1-GFP (green, mitochondrial marker) and labeled with HMSiR$_{680}$ (magenta). Contrast in the magenta channel has been adjusted to clearly show off-target signal. Overlapping signal appears white. Scale bar = 10 µm.

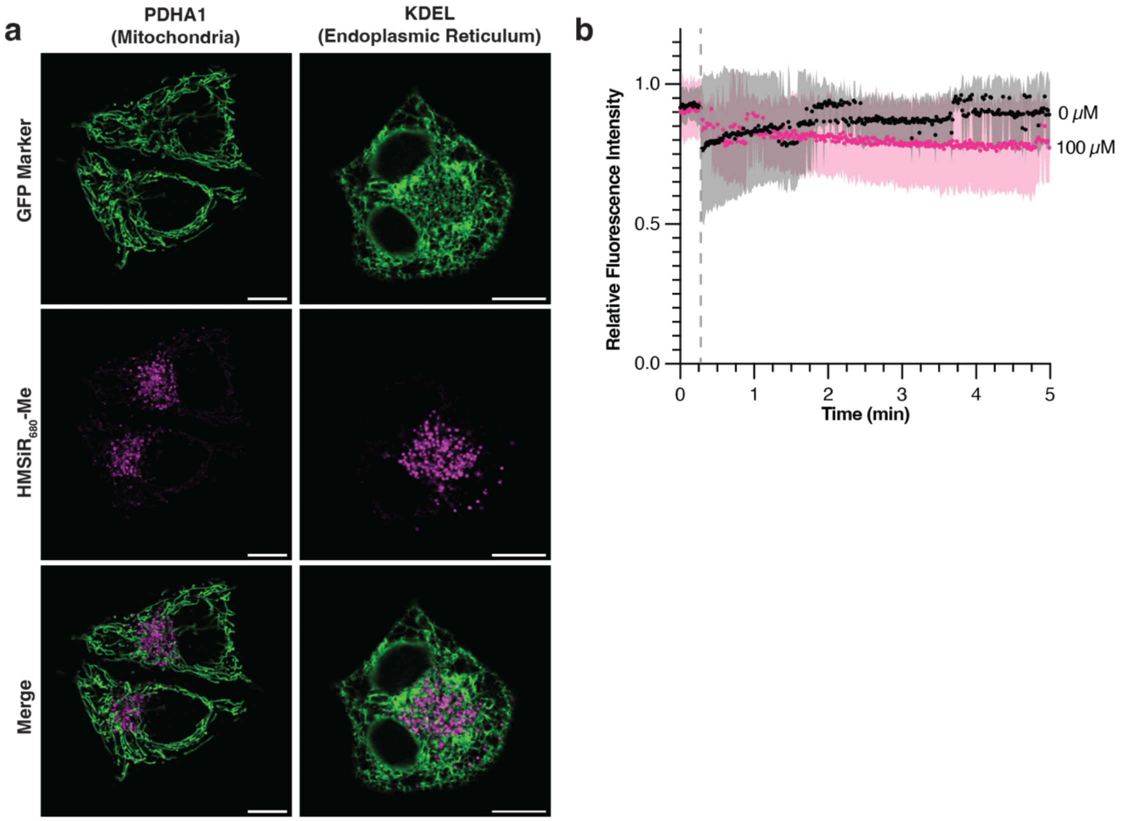

**Figure S2.** HMSiR$_{680}$-Me selectively targets acidic vesicles in live HeLa cells. (a) Representative colocalization between HMSiR$_{680}$-Me (magenta) and GFP organelle markers for mitochondria and the ER (green). Regions of the cell in which the magenta and green signals overlap appear white. Scale bars = 10 μm. (b) Fluorescence intensity of GFP in HeLa cells labeled with LAMP1-GFP and incubated with the indicated concentration of chloroquine diphosphate (CQ) for 5 min. CQ solution was introduced to the cells 20 s into the 5 min imaging period (represented by dashed line). Data plotted in panel (b) represents 3 biological replicates; shaded area represents standard deviation.

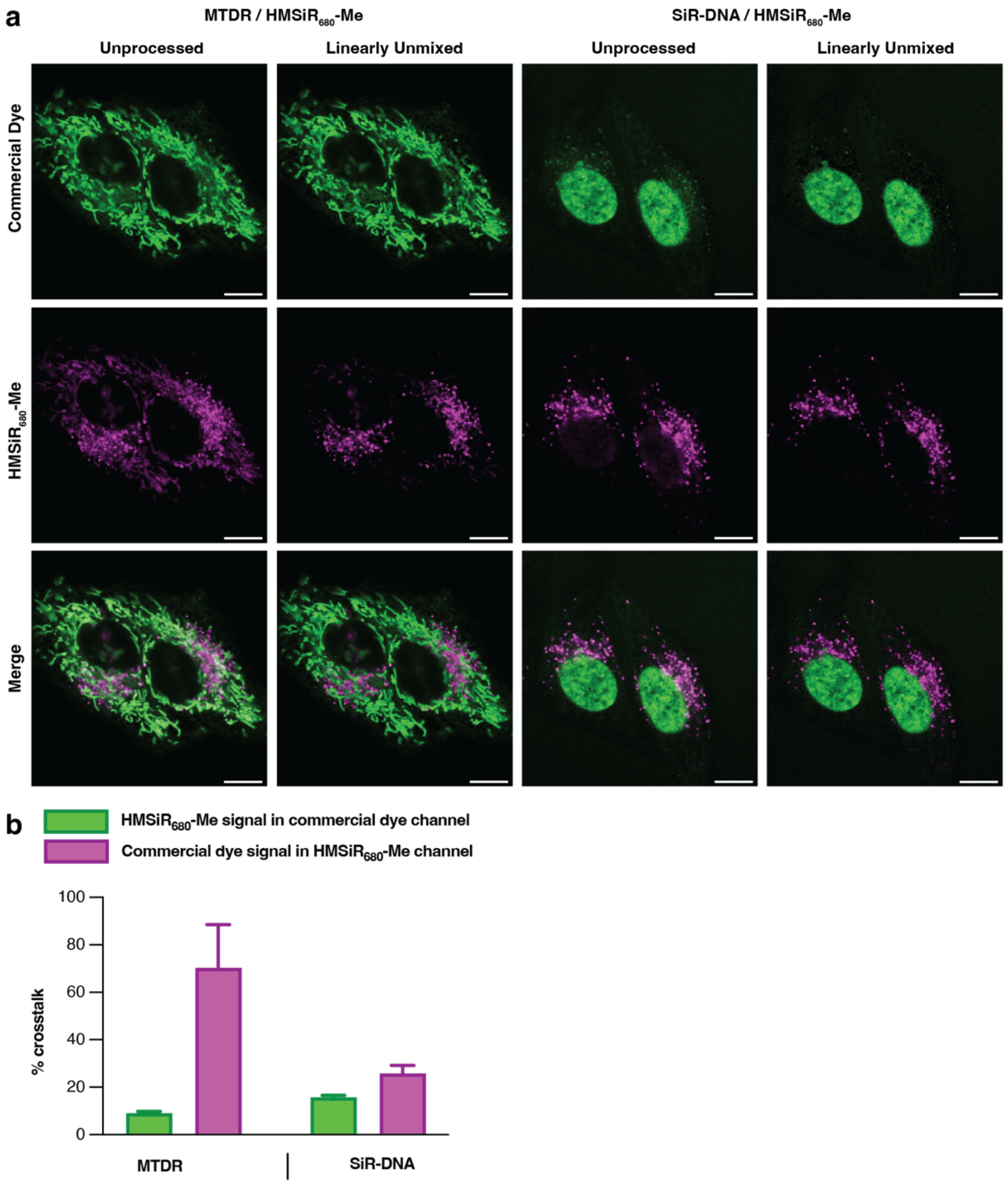

**Figure S3.** HMSiR$_{680}$-Me can be used alongside other far-red dyes in two-color imaging experiments. (a) HeLa cells labeled with MitoTracker Deep Red (MTDR) or SiR-DNA (green) and HMSiR$_{680}$-Me (magenta). Unprocessed images and images that have been linearly unmixed with the Leica Stellaris Dye Separation tool are shown. Scale bars = 10 µm. (b) Percent crosstalk for each unprocessed spectral channel in (a). Crosstalk values were calculated as described in the supplementary methods. Data represents 8 technical replicates; error bars represent standard deviation. Dye concentrations were as follows: 500 nM HMSiR$_{680}$-Me, 50 nM MitoTracker Deep Red, 1 µM SiR-DNA.

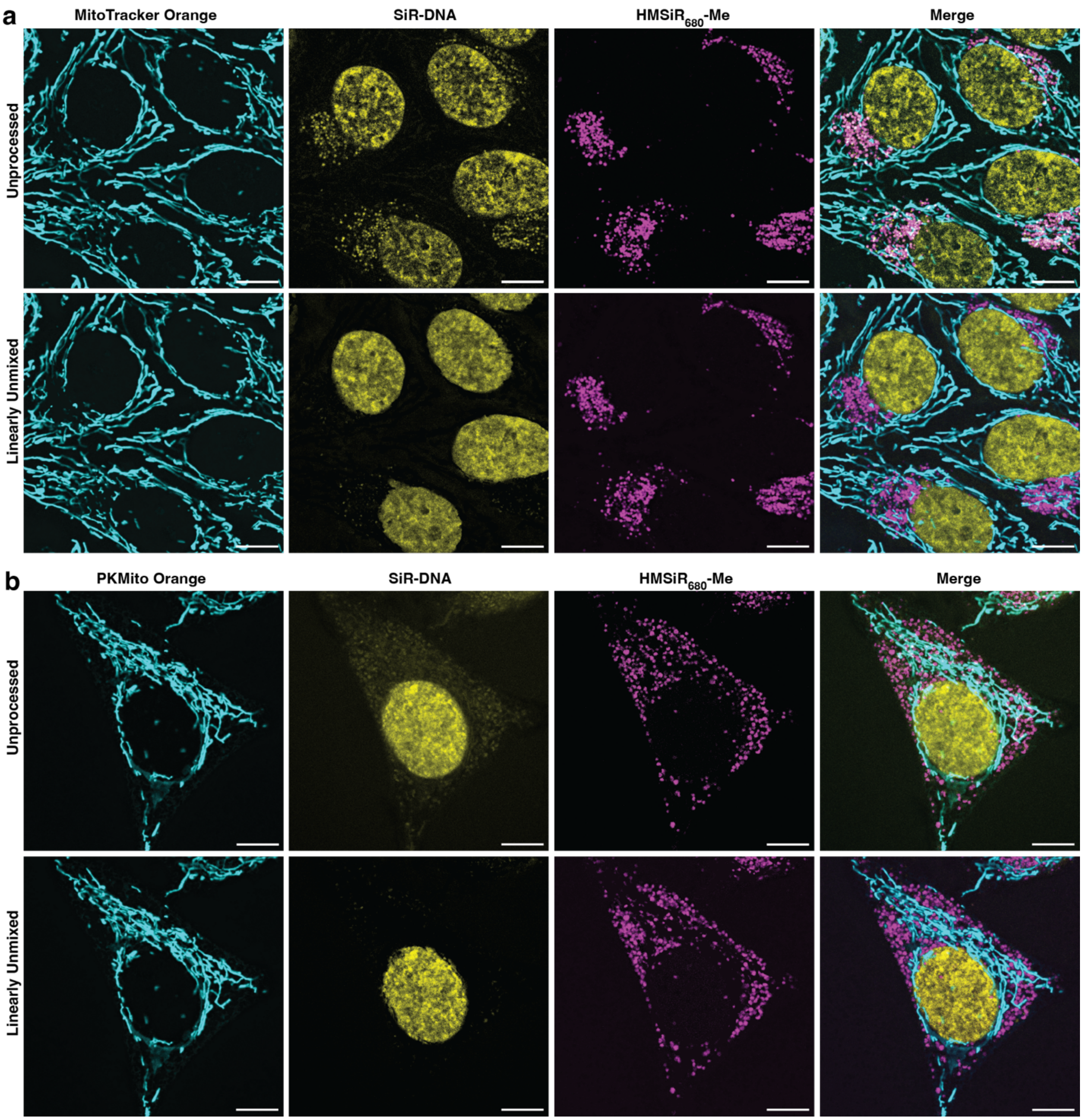

**Figure S4.** HMSiR$_{680}$-Me can be used alongside other red and far-red dyes in three-color imaging experiments. HeLa cells labeled with (a) MitoTracker Orange or (b) PKMito Orange (cyan), SiR-DNA (yellow), and HMSiR$_{680}$-Me (magenta). Unprocessed images and images that have been linearly unmixed with the Leica Stellaris Dye Separation tool are shown. Scale bars = 10 µm. Dye concentrations were as follows: 500 nM HMSiR$_{680}$-Me, 50 nM MitoTracker Deep Red, 1 µM SiR-DNA, 100 nM MitoTracker Orange, 1x PKMito Orange.

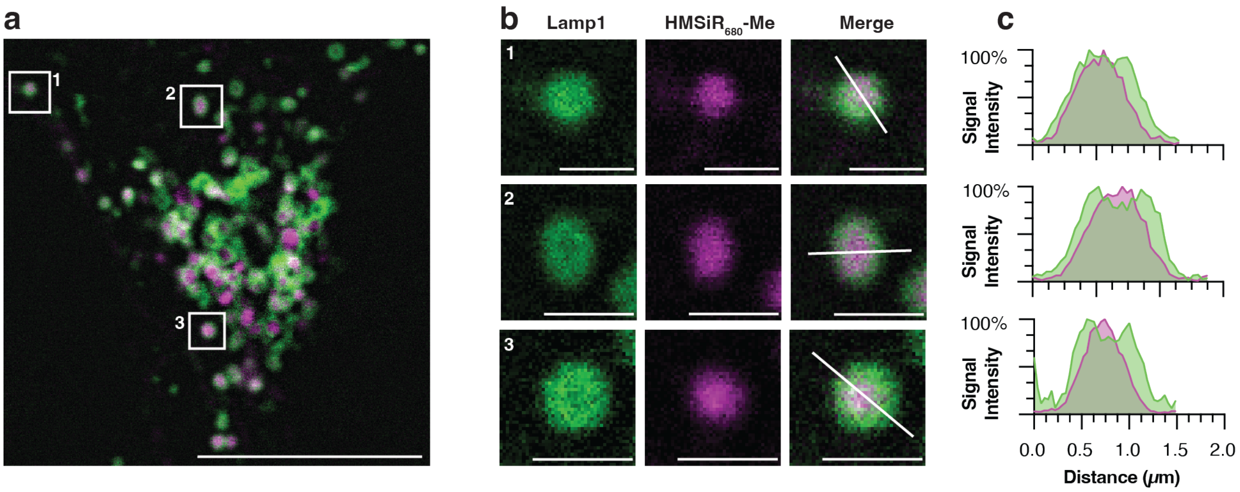

**Figure S5**. Confocal microscopy does not effectively resolve HMSiR$_{680}$-Me localization to the lysosomal lumen in HeLa cells. (a) A HeLa cell labeled with HMSiR$_{680}$-Me (magenta) and LAMP1-HaloTag and SiR-chloroalkane (green) and imaged via confocal microscopy. Scale bar = 10 µm. (b) Insets from part (a). Scale bars = 1 µm. (c) Line profile diagrams along the lines shown in part (b). Maximum intensity for each channel was scaled. Dye concentrations were 500 nM for HMSiR$_{680}$-Me and 2 µM for SiR-chloroalkane.

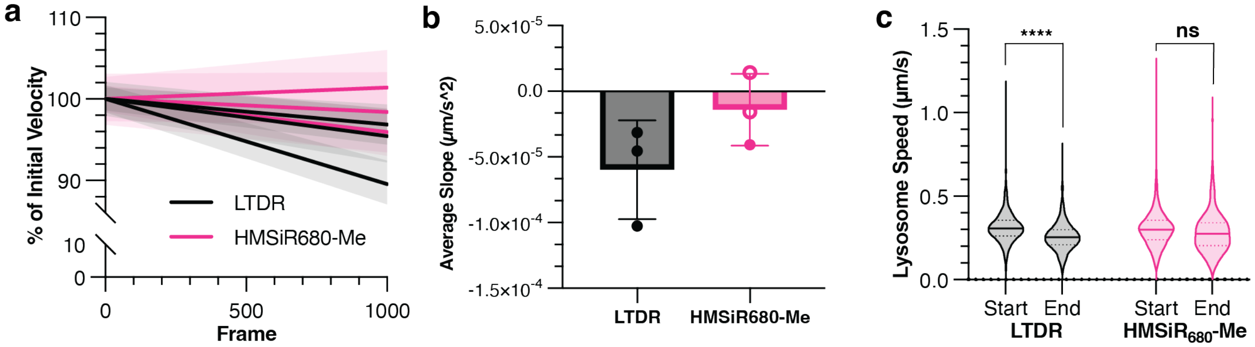

**Figure S6**. HMSiR$_{680}$-Me does not affect lysosome motility. (a) Linear regressions for lysosome speed over time for HeLa cells labeled with 50 nM LTDR or 500 nM HMSiR$_{680}$-Me. Each slope represents a biological replicate. Initial velocity was normalized, and 95% confidence intervals are shown. (b) Average deceleration of lysosomes for the data shown in (a). Filled-in data points represent slopes that are significantly nonzero ($p < 0.0001$, F test); empty points indicate $p \geq 0.0840$. (c) Violin plots showing tracks beginning in the first 50 (“start”) or last 50 (“end”) frames of the data plotted in (a). Each plot in panel (c) represents at least 171 tracks across 3 biological replicates. “****” indicates $p < 0.0001$; “ns” indicates $p > 0.05$ (unpaired $t$ test with Welch’s correction).

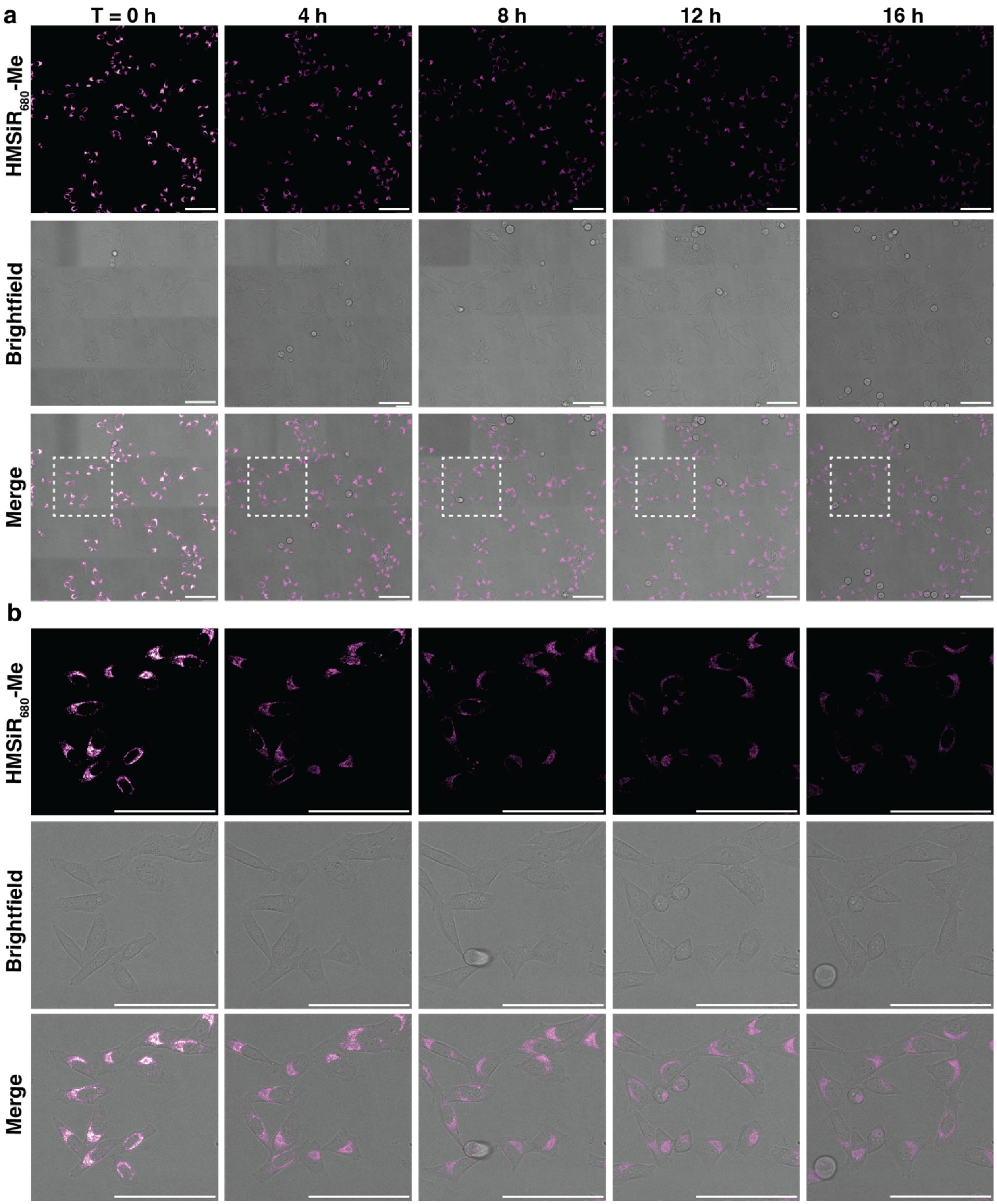

**Figure S7**. HMSiR$_{680}$-Me can label live cells for long time lapse confocal microscopy. (a) Confocal time lapse images of HeLa cells labeled with 500 nM HMSiR$_{680}$-Me (magenta) over 16 h. (b) Inset from part (a). Scale bars = 20 μm.

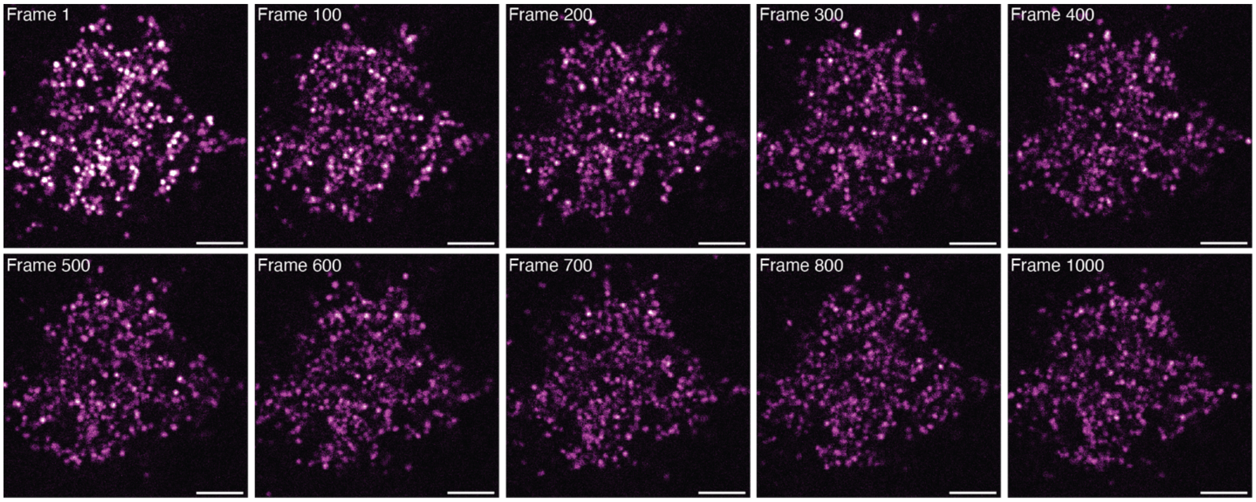

**Figure S8.** STED time lapse images of HeLa cells labeled with 1 µM $HMSiR_{680}$-Me with a frame rate of 1.5 fps and pixel size of 50 nm. Scale bars = 2 µm.

**II. Supplementary Video Descriptions**

**Movie S1:** $HMSiR_{680}$-Me is pH-dependent in live cells.
HeLa cells labeled with $HMSiR_{680}$-Me (magenta) or transfected with LAMP1-GFP (green) were imaged at 1 fps. After 20 frames, the imaging media was quickly replaced with media containing the indicated concentration of chloroquine diphosphate (CQ). Video frame rate is 36 fps. Scale bars = 200 µm.

**Movie S2:** $HMSiR_{680}$-Me can be used with other red and far-red dyes in multicolor imaging experiments.
HeLa cell labeled with $HMSiR_{680}$-Me (magenta), PKMO (cyan), and SiR-DNA (yellow). Dyes were linearly unmixed using the Leica Stellaris Dye Separation tool. Video frame rate is 24 fps. Scale bar = 10 µm.

**Movie S3:** $HMSiR_{680}$-Me photobleaches slowly and does not affect cell health.
HeLa cells labeled with either $HMSiR_{680}$-Me or LTDR at the indicated concentration were excited at the indicated laser power and imaged for 1 h. Videos are representative of n = 3 replicates for each condition. Video frame rate is 200 fps. Scale bars = 20 µm.

**Movie S4:** $HMSiR_{680}$-Me does not affect lysosomal motility.
HeLa cells labeled with either 500 nM $HMSiR_{680}$-Me or 50 nM LTDR were imaged at 1fps. Video frame rate is 100 fps. Scale bars = 20 µm.

**Movie S5:** $HMSiR_{680}$-Me is compatible with overnight confocal imaging.
HeLa cells were labeled with $HMSiR_{680}$-Me, and a region of 4x4 frames was imaged every 2 min for 16 h and stitched together. The marked region is enlarged on the right to show finer detail. Focal plane was maintained using Leica's Adaptive Focus Control system. Video frame rate is 24 fps. Scale bar = 100 µm.

**Movie S6:** $HMSiR_{680}$-Me is compatible with timelapse STED imaging with excellent temporal resolution.
A HeLa cell labeled with $HMSiR_{680}$-Me was imaged via STED at 1.5 fps with a pixel size of 50 nm. Video frame rate is 24 fps. Scale bar = 2 µm.

**Movie S7:** $HMSiR_{680}$-Me can be used with SiR in a two-color multiplexed imaging experiment.
HeLa cell labeled with MAO-$N_3$/SiR-DBCO (green) and $HMSiR_{680}$-Me (magenta). Dyes were simultaneously excited at 645 nm and emission was detected across three spectral windows between 655 nm and 766 nm. Dyes were linearly unmixed using the Leica Stellaris Dye Separation tool. Video frame rate is 24 fps. Scale bar = 5 µm.

**Movie S8:** $HMSiR_{680}$-Me can be used with SiR in a two-color multiplexed STED imaging experiment.
HeLa cell labeled with MAO-$N_3$/SiR-DBCO (green) and $HMSiR_{680}$-Me (magenta). Dyes were simultaneously excited at 645 nm and emission was detected across two spectral windows between 655 nm and 766 nm. Dyes were linearly unmixed using the Leica Stellaris Dye Separation tool. Video frame rate is 12 fps. Scale bar = 2 µm.

### III. Supplementary Methods

#### A. In Vitro Spectroscopy

##### 1. Determination of p$K_{cycl}$

Samples of each fluorophore were prepared at 15 μM in 0.2 M phosphate buffer at half-integral pH values between 1.5 and 10, and 300 μL was pipetted in triplicate into a black 96-well plate with a clear bottom. Control wells for each pH value were also prepared. Using a BioTek Synergy H1 plate reader, the absorbance for each well was measured at its maximum absorbance wavelength. The absorbance values from the control wells were subtracted from the measured absorbance, and the data were normalized such that the maximum absorbance measurement for each dye was equal to 1.0. The data were then plotted using GraphPad Prism 9.0 and sigmoidal curve fits were performed. p$K_{cycl}$ was set as the pH at which absorbance is one-half of the maximum.

##### 2. Photobleaching Measurements

500 nM solutions of each fluorophore were prepared in 0.2 M phosphate buffer at a pH of 4.5, and 300 μL was pipetted in triplicate into a black 96-well plate with a clear bottom. Using a BioTek Synergy H1 plate reader, wells were excited at 650 nm and emission was detected at 680 nm, and 695 scans were performed on each well at a scan interval of 11s. The data were normalized such that the maximum fluorescence intensity measurement for each dye was equal to 1.0.

#### B. Microscopy

##### 1. Materials and General Information

HeLa cells were cultured in Dulbecco's Modified Eagle Medium (DMEM, 4.5 g/L glucose) supplemented with 10% (v/v) fetal bovine serum (FBS) and 1% Pen Strep in a 37˚C humidified incubator at 5% $CO_2$. Cells were split every 2-4 days and discarded upon reaching passage 12. Unless otherwise noted, cells were plated on #1.5H two-, four-, or eight-well glass bottom μ-slides (Ibidi) 16-48 hours prior to imaging. All GFP transfection was done using CellLight BacMam GFP reagents (ThermoFisher). CMV-LAMP1-HaloTag was a gift from Jennifer Lippincott-Schwartz (Addgene plasmid # 164209; http://n2t.net/addgene:164209; RRID:Addgene_164209), and transfection was done using the FuGENE HD transfection system (Promega). MitoTracker Deep Red (MTDR), MitoTracker Orange, and LysoTracker Deep Red (LTDR) were purchased from ThermoFisher. SiR-DNA and PKmito Orange (PKMO) were purchased from Spirochrome.

Microscopy was performed on a Leica STELLARIS 8 microscope (Leica Microsystems) equipped with a HC Plan-Apo 10x/0.4NA dry objective, a HC Plan-Apo 63x/1.4NA water immersion objective with a motorized correction collar, a HC Plan-Apo 63x/1.4NA oil immersion objective, a HC Plan-Apo 100x/1.4NA oil immersion objective, a pulsed white-light laser (440 nm- 790 nm; 440 nm: > 1.1 mW; 488 nm: > 1.6 mW; 560 nm: > 2.0 mW; 630 nm: > 2.6 mW; 790 nm: > 3.5 mW, 78 MHz), a pulsed 775 nm STED laser, a Leica DMi8 CS scanhead, and spectral detectors with sliding emission windows. When one or two detectors was required, imaging was performed using only HyD X detectors in counting mode. When three detectors were required, a single HyD S detector was also used. Live cell imaging conditions (37˚C, humidified 5% $CO_2$) were maintained in a blacked out cage enclosure from Okolab.

### 2. Supplementary Table 1: Confocal and STED Imaging Parameters

| Figure | Compound | Concentration | Excitation | | Depletion | | Detection | Objective Lens | Pixel | Dwell | Image | Frame | Frames | Multicolor |
|---|---|---|---|---|---|---|---|---|---|---|---|---|---|---|
| | | | Laser (nm) | Power (%) | Laser (nm) | Power (%) | Window | | Size (nm) | Time (µs) | Size (px) | Rate (fps) | | Acquisition Mode |
| 1d | HMSiR680 | 2 µM | 680 | 2 | - | - | 691-829 | 63x water | 180 | 3.8375 | 1024 x 1024 | - | 1 | - |
| 1e | HMSiR680-Me | 2 µM | 680 | 2 | - | - | 691-829 | 63x water | 180 | 3.8375 | 1024 x 1024 | - | 1 | - |
| | GFP | - | 489 | 20 | - | - | 500-642 | | | | | | | |
| S1b | HMSiR680 | 500 nM | 680 | 2 | - | - | 691-807 | 63x water | 76 | 3.8375 | 1024 x 1024 | - | 1 | line sequential |
| | GFP | - | 489 | 15-25 | - | - | 500-642 | | | | | | | |
| 2a/S2a | HMSiR680-Me | 500 nM | 680 | 10 | - | - | 691-829 | 63x oil | 40-140 | 3.8375 | 1024 x 1024 | - | 1 | line sequential |
| | MTDR | 50 nM | 641 | 2 | - | - | 655-800 | | | | | | | |
| 3a/S3a | HMSiR680-Me | 500 nM | 695 | 20 | - | - | 705-829 | 63x oil | 60 | 0.82625 | 1024 x 1024 | - | 1 | frame sequential |
| | SiR-DNA | 1 µM | 650 | 20 | - | - | 662-700 | | | | | | | |
| 3b/S3a | HMSiR680-Me | 500 nM | 680 | 20 | - | - | 690-829 | 63x oil | 120 | 1.725 | 512 x 512 | - | 1 | frame sequential |
| | MT Orange | 100 nM | 560 | 1 | - | - | 570-620 | | | | | | | |
| | SiR-DNA | 1 µM | 650 | 20 | - | - | 662-700 | | | | | | | |
| 3c/S4a | HMSiR680-Me | 500 nM | 680 | 20 | - | - | 690-829 | 63x oil | 60 | 1.575 | 1024 x 1024 | - | 1 | frame sequential |
| | PKMO | 1x | 591 | 2 | - | - | 603-653 | | | | | | | |
| | SiR-DNA | 1 µM | 650 | 20 | - | - | 662-700 | | | | | | | |
| 3d/S4b | HMSiR680-Me | 500 nM | 680 | 20 | - | - | 690-829 | 63x oil | 60 | 1.575 | 1024 x 1024 | - | 1 | frame sequential |
| | SiR-chloroalkane | 2 µM | 645 | 10 | 775 | 20 | 655-670 | | | | | | | |
| 4 | HMSiR680-Me | 500 nM | 680 | 30 | 775 | 10 | 695-765 | 100x oil | 37 | 7.6875 | 512 x 512 | - | 1 | line sequential |
| | SiR-chloroalkane | 2 µM | 645 | 7 | - | - | 655-670 | | | | | | | |
| S5 | HMSiR680-Me | 500 nM | 680 | 25 | - | - | 695-765 | 100x oil | 37 | 7.6875 | 512 x 512 | - | 1 | line sequential |
| 5c | HMSiR680-Me | 500 nM | 680 | 20 | - | - | 690-845 | 63x oil | 144 | 2.2625 | 512 x 512 | 0.96 | 3465 | - |
| 5d | LTDR | 500 nM | 680 | 5 | - | - | 690-845 | 63x oil | 144 | 2.2626 | 512 x 512 | 0.96 | 3465 | - |
| 5e | LTDR | 50 nM | 680 | 20 | - | - | 690-845 | 63x oil | 144 | 2.2625 | 512 x 512 | 0.96 | 3465 | - |
| S7 | HMSiR680-Me | 500 nM | 684 | 10 | - | - | 695-757 | 63x oil | 180 | 3.8375 | 3798 x 3828 | 0.00208 | 408 | - |
| S8 | HMSiR680-Me | 1 µM | 680 | 20 | 775 | 15 | 690-765 | 100x oil | 51 | 2.0875 | 512 x 512 | 1.54 | 1000 | - |
| | SiR-DBCO | 100 nM | | | | | 655-673 | | | | | | | |
| | - | - | | | | | 673-695 | | | | | | | |
| 6a | HMSiR680-Me | 500 nM | 645 | 20 | - | - | 695-766 | 100x oil | 59 | 15.3875 | 1024 x 1024 | - | 1 | simultaneous |
| | SiR-DBCO | 100 nM | | | | | 655-673 | | | | | | | |
| | - | - | | | | | 673-695 | | | | | | | |
| 6b | HMSiR680-Me | 500 nM | 645 | 20 | - | - | 695-766 | 100x oil | 101 | 7.6875 | 512 x 512 | 0.387 | 300 | simultaneous |
| | SiR-DBCO | 100 nM | | | | | 655-680 | | | | | | | |
| 6c | HMSiR680-Me | 1 µM | 645 | 20 | 775 | 20 | 686-766 | 100x oil | 10 | 7.6875 | 1024 x 1024 | - | 1 | simultaneous |
| | SiR-DBCO | 100 nM | | | | | 655-680 | | | | | | | |
| 6d | HMSiR680-Me | 1 µM | 645 | 20 | 775 | 20 | 686-766 | 100x oil | 31 | 8.35 | 1024 x 256 | 0.385 | 60 | simultaneous |

| Movie | Compound | Concentration | Excitation | | Depletion | | Detection | Lens | Pixel | Dwell | Image | Frame | Frames | Multicolor |
|---|---|---|---|---|---|---|---|---|---|---|---|---|---|---|
| | | | Laser (nm) | Power (%) | Laser (nm) | Power (%) | Window | | Size (nm) | Time (µs) | Size (px) | Rate (fps) | | Acquisition Mode |
| Movie S1 | HMSiR680-Me | 2 µM | 680 | 10 | - | - | 690-750 | 10x dry | 2275 | 1.725 | 512 x 512 | 1.15 | 364 | - |
| | GFP | - | 473 | 20 | - | - | 483-750 | 10x dry | 3033 | 1.725 | 512 x 512 | 1.19 | 378 | - |
| | PKMO | 1x | 591 | 2 | - | - | 603-653 | | | | | | | |
| | SiR-DNA | 1 µM | 650 | 20 | - | - | 662-700 | | | | | | | |
| Movie S2 | HMSiR680-Me | 500 nM | 680 | 20 | - | - | 690-829 | 63x oil | 60 | 1.575 | 1024 x 1024 | 0.074 | 300 | frame sequential |
| | HMSiR680-Me | 500 nM | 680 | 20 | - | - | 690-845 | 63x oil | 144 | 2.2625 | 512 x 512 | 0.96 | 3465 | - |
| | LTDR | 500 nM | 680 | 5 | - | - | 690-845 | 63x oil | 144 | 2.2626 | 512 x 512 | 0.96 | 3465 | - |
| Movie S3 | LTDR | 50 nM | 680 | 20 | - | - | 690-845 | 63x oil | 144 | 2.2625 | 512 x 512 | 0.96 | 3465 | - |
| Movie S4 | HMSiR680-Me | 500 nM | 680 | 20 | - | - | 690-845 | 63x oil | 144 | 2.2625 | 512 x 512 | 0.96 | 1000 | - |
| | LTDR | 50 nM | 680 | 20 | - | - | 690-845 | 63x oil | 144 | 2.2625 | 512 x 512 | 0.96 | 1000 | - |
| Movie S5 | HMSiR680-Me | 500 nM | 684 | 10 | - | - | 695-757 | 63x oil | 180 | 3.8375 | 3798 x 3828 | 0.00833 | 408 | - |
| Movie S6 | HMSiR680-Me | 1 µM | 680 | 20 | 775 | 15 | 690-765 | 100x oil | 51 | 2.0875 | 512 x 512 | 1.54 | 1000 | - |
| | SiR-DBCO | 100 nM | | | | | 655-673 | | | | | | | |
| | - | - | | | | | 673-695 | | | | | | | |
| Movie S7 | HMSiR680-Me | 500 nM | 645 | 20 | - | - | 695-766 | 100x oil | 101 | 7.6875 | 512 x 512 | 0.387 | 300 | simultaneous |
| Movie S8 | SiR-DBCO | 100 nM | 645 | 20 | 775 | 20 | 655-680 | 100x oil | 31 | 8.35 | 1024 x 256 | 0.385 | 60 | simultaneous |
| | HMSiR680-Me | 1 µM | | | | | 686-766 | | | | | | | |

**3. General Labeling Protocol**

At least 16 hours after plating, cells were washed 3X with warm DPBS, then treated with a solution of the small-molecule fluorophore(s) in pre-warmed OptiMEM ph(-). Cells were incubated at 37˚C and 5% $CO_2$ for 30 min, washed once with warm DMEM ph(-), and imaged in DMEM ph(-).

**4. Colocalization and Determination of Pearson's Correlation Coefficients**

The day before imaging, cells were plated on 8-well plates at a density of 1.5 x $10^4$ cells per well. After one hour, 15 μL of a CellLight BacMam GFP reagent for organelles of interest (lysosome, late endosome, early endosome, mitochondria, or ER) was added to each well and the media was gently mixed with the 15 μL pipette. The cells were further incubated for 12-20 hours. On the day of imaging, cells were labeled with 500 nM $HMSiR_{680}$-Me or $HMSiR_{680}$ as described above, and 12 images per well were obtained. For each image, transfected cells were manually identified in CellProfiler[1] and Pearson's correlation coefficients for overlap of GFP/$HMSiR_{680}$-Me signal within each cell was measured using the "MeasureColocalization" function with a threshold of 15%.

**5. Imaging with Chloroquine**

Cells were plated on 8-well μ-slides with ibiTreat polymer coverslips (Ibidi) at a density of 1.5 x $10^4$ cells per well. After one hour, 15 μL of CellLight BacMam Lamp1-GFP reagent (ThermoFisher) was added to control wells and the media was gently mixed 5 times with the 15 μL pipette. The cells were further incubated for 16 hours. On the day of imaging, all wells were washed 3x with warm DPBS. In the control wells (those labeled with Lamp1-GFP), the media was immediately exchanged with 300 μL warm DMEM(ph-). The remaining wells were labeled with 2 mM $HMSiR_{680}$-Me as described above. Each well was imaged for 20 frames, then image acquisition was stopped and without disturbing the slide, the media was quickly replaced with 300 μL DMEM(ph-) containing chloroquine diphosphate. Image acquisition was restarted immediately and continued for five minutes. Each condition was performed in triplicate. Frames from before and after media exchange were concatenated in FIJI, and fluorescence intensity was measured for each frame within the default thresholded region. Data were normalized such that the maximum fluorescence intensity for each condition was set equal to 1.0.

**6. Calculating Crosstalk**

Cells were labeled with either $HMSiR_{680}$-Me, MTDR, or SiR-DNA then imaged according to the 2-color parameters given in Supplementary Table 2. To calculate the % crosstalk in the $HMSiR_{680}$-Me channel, cells labeled only with the commercial dye (MTDR or SiR-DNA) were imaged in both spectral channels. Each channel was automatically thresholded and the average signal intensity of the thresholded region was measured. The % crosstalk in the $HMSiR_{680}$-Me channel was obtained for cells labeled with only commercial dye by dividing the signal intensity in the $HMSiR_{680}$-Me channel by the signal intensity in the commercial dye channel. Similarly, to determine the % crosstalk in the commercial dye channel, cells labeled only with $HMSiR_{680}$-Me were imaged in both channels, and the % crosstalk was calculated by dividing the signal intensity in the commercial dye channel by the signal intensity in the $HMSiR_{680}$-Me channel.

**7. Two-Color Imaging with HMSiR$_{680}$-Me and SiR**

*MAO-N$_3$/SiR-DBCO and HMSiR$_{680}$-Me*

Cells were plated on 2-well µ-slides at a density of 4.7 x $10^4$ cells per well 36-48 hours before imaging. On the day of imaging, cells were washed 3x with warm DPBS, then labeled with 1.5 mL of 200 nM MAO-N$_3$ in pre-warmed OptiMEM(ph-) and incubated for 1h. They were washed once with DMEM(ph-) and labeled with 1.5 mL of 100 nM SiR-DBCO in prewarmed OptiMEM(ph-) for 1h. The media was then exchanged with DMEM(ph-) with 10% FBS and incubated for 1h. The cells were washed once with DMEM(-ph) and labeled with 500 nM HMSiR$_{680}$-Me in prewarmed OptiMEM(ph-) for 30 min. Finally, the cells were washed once with DMEM(-ph) and imaged in 1.5 mL DMEM(ph-).

*Lamp1-Halo/SiR-chloroalkane and HMSiR$_{680}$-Me*

2 days before imaging, cells were plated on 4-well µ-slides at a density of 2.2 x $10^4$ cells per well. The next day, each well was transfected with a mixture of 60 µL OptiMEM, 500 ng of plasmid encoding Lamp1-HaloTag, and 1.5 µL room temperature FuGENE HD reagent (Promega). Cells were incubated for 6h, then media was replaced with fresh warm DMEM including 10% v/v FBS and 1% v/v Penn Strep. On the day of imaging, cells were washed 3x with warm DPBS then labeled with a solution of 2 µM SiR-chloroalkane and 500 nM HMSiR$_{680}$-Me in prewarmed OptiMEM. Cells were incubated for 1h, then washed 3x with DMEM(ph-). The media was replaced with fresh DMEM(ph-) and cells were imaged.

**8. Photobleaching Measurements in Cellulo**

Cells were labeled with either LysoTracker Deep Red (ThermoFisher) or HMSiR$_{680}$-Me and imaged as described above. A background subtraction was performed and fluorescence intensity was measured for each frame within the default thresholded region. The data were normalized such that the maximum fluorescence intensity measurement for each dye was equal to 1.0.

**9. Lysosome Velocity Measurements**

Cells were labeled with either LysoTracker Deep Red (ThermoFisher) or HMSiR$_{680}$-Me and imaged as described above. A sliding parabola background subtraction with a pixel size of 7 was performed. The TrackMate FIJI plugin[2] was used to track particle movement. The LoG detector was used with an estimated blob diameter of 0.4 µm and a threshold of 2, and the Simple LAP tracker was used with a linking max distance of 2 µm, a gap-closing max distance of 3 µm, and a gap-closing max frame gap of 2. A track filter was set to discard tracks with a mean velocity above 1.5 µm/s, and the track quality filter was adjusted manually to retain approximately 1000-2000 tracks per cell. Average Track Velocity vs. Track Start Time was plotted, the data were exported to GraphPad Prism 9, and linear regressions were calculated for each replicate.

**C. Chemical Synthesis**

**1. Materials and General Information**

Chemicals used for synthesis were prepared according to the literature or purchased from commercial sources and used without purification. THF used in the preparation of (**3**) and (**4**) was dried over 4Å molecular sieves before use. Preparative flash chromatography was performed on a Teledyne Isco CombiFlash Rf using pre-packed RediSep gold or silver columns and the solvent gradients indicated. Preparative HPLC was performed on a Waters 150 HPLC using a 95-10% Water/Acetonitrile gradient with 0.1% TFA additive. The HPLC was equipped with a reverse-phase C18 column, a 200-800 nm UV/vis detector, an automatic fraction collector, and an autosampler with a 7 mL injection loop. Low-resolution mass spectra were measured on a Waters SQD2 using a 90-10% Water/Acetonitrile gradient with a 0.1% formic acid additive. The LCMS was equipped with a reverse-phase C18 column, a 200-800 nm UV/vis detector, a 300-1000 nm fluorescence detector, and an autosampler. NMR spectra were measured on a Bruker NEO500 spectrometer (500 MHz for $^{1}H$ and 126 MHz for $^{13}C$) or a Bruker AV600 spectrometer (600 MHz for $^{1}H$ and 151 MHz for $^{13}C$) and analyzed with Mestrelab MestReNova 14.2. Chemical shift values are reported in ppm in reference to residual solvent peaks of $CDCl_3$ (7.26 for $^{1}H$, 77.16 for $^{13}C$) or $CD_3OD$ (3.31 for $^{1}H$, 49.00 for $^{13}C$). Coupling constants (*J*) are reported in Hz. Procedures for the synthesis of silicon rhodamine scaffolds were adapted from a recent report.[3]

### 2. Synthesis of HMSiR$_{680}$ and HMSiR$_{680}$-Me

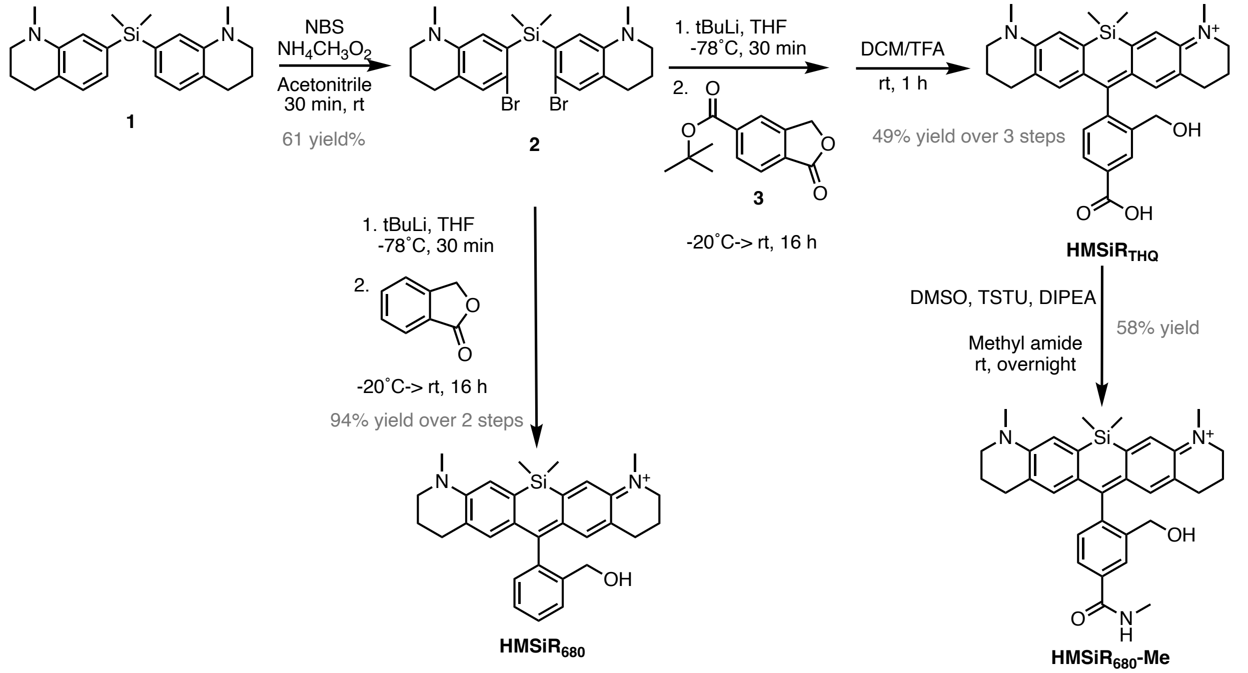

**bis(6-bromo-1-methyl-1,2,3,4-tetrahydroquinolin-7-yl)dimethylsilane** (**2**): Dimethylbis(1- methyl-1,2,3,4- tetrahydroquinolin-7-yl)silane (**1**) was prepared according to a previous report[4] and the following procedure was adapted from the literature.[5] In a 20 mL scintillation vial, **1** (152 mg, 0.434 mmol) and ammonium acetate (7 mg, 0.087 mmol, 0.2 eq) were dissolved in acetonitrile (2.89 mL) and cooled to 0˚C. The solution was stirred under vacuum for 20 minutes to degas and then charged with $N_2$. N-bromosuccinimide (NBS, 158 mg, 0.889 mmol, 2.05 eq) was added at once and the vial was resealed, then the reaction was stirred under nitrogen and allowed to warm to room temperature over 30 min. The acetonitrile was removed, the crude reaction mixture was dissolved in $CH_2Cl_2$ and washed with brine. The aqueous layer was extracted twice more with $CH_2Cl_2$, and the combined organic fractions were dried over $Na_2SO_4$. The solvent was evaporated and the crude product was purified by flash chromatography (0%-30% ethyl acetate in hexanes) to yield a pink solid (102 mg, 61% yield). $^{1}H$ NMR (500 MHz, $CDCl_3$) δ 7.09 (s, 2H), 6.68 (s, 2H), 3.18 (t, $J$ = 5.7 Hz, 4H), 2.79 (s, 6H), 2.72 (t, $J$ = 6.4 Hz, 4H), 1.95 (p, $J$ = 6.3 Hz, 4H), 0.72 (s, 6H). $^{13}C$ NMR (126 MHz, $CDCl_3$) δ 145.3, 136.5, 132.6, 126.3, 120.2, 116.4, 39.2, 27.6, 22.3, -0.65. LRMS calc'd for $C_{22}H_{29}Br_2N_2Si$ $[M+H]^+$ *m/z* 507.0467, found 507.1824.

**$HMSiR_{680}$**: In a flame-dried flask, **2** (34 mg, 0.067 mmol) was dissolved in anhydrous THF (1.29 mL) and cooled to -78˚C under $N_2$. *tert*-butyllithium (1.7 M in pentane, 173 µL, 0.294 mmol, 4.4 eq) was added dropwise and the reaction was allowed to stir for 30 min. In a separate flame-dried flask, phthalide (20 mg, 0.147 mmol, 2.2 eq) was dissolved in anhydrous THF (1.29 mL). The reaction was warmed to -20˚C and the phthalide solution was added dropwise over 30 min. The reaction was allowed to slowly warm to room temperature overnight, and acetic acid (0.5 mL) was added to yield an immediate blue color. The solvent was removed and the crude residue was dissolved in $CH_2Cl_2$ and washed with brine. The aqueous layer was extracted twice more with $CH_2Cl_2$, and the combined organic fractions were dried over $Na_2SO_4$. The solvent was evaporated and the crude product was purified by flash chromatography (0%-50% ethyl acetate in hexanes) to yield a blue solid (29 mg, 94% yield). $^{1}$H NMR (600 MHz, MeOD) δ 7.72 (d, *J* = 7.7 Hz, 1H), 7.58 (t, *J* = 7.4 Hz, 1H), 7.45 (t, *J* = 7.4 Hz, 1H), 7.22 (s, 2H), 7.10 (d, *J* = 7.4 Hz, 1H), 6.70 (s, 2H), 4.32 (s, 2H), 3.60 (t, *J* = 5.5 Hz, 4H), 3.33 (s, 6H), 2.51 – 2.45 (m, 4H), 1.91 (p, *J* = 6.1 Hz, 4H), 0.57 (d, *J* = 8.3 Hz, 6H). $^{13}$C NMR (151 MHz, MeOD) δ 167.7, 153.4, 148.5, 140.6, 139.2, 138.6, 130.2, 130.0, 128.9, 128.1, 128.0, 126.0, 121.0, 62.3, 53.5, 39.8, 28.3, 22.1, -1.0, -1.3. LRMS calc'd for $C_{30}H_{35}N_2OSi^+$ $[M]^+$ *m/z* 467.2513, found 467.4381.

**$HMSiR_{THQ}$**: In a flame-dried flask, **2** (160 mg, 0.315 mmol) was dissolved in anhydrous THF (6 mL) and cooled to -78˚C under $N_2$. *tert*-butyllithium (1.7 M in pentane, 0.815 mL, 1.385 mmol, 4.4 eq) was added dropwise and the reaction was allowed to stir for 30 min. In a separate flame-dried flask, **3** (162 mg, 0.692 mmol, 2.2 eq) was dissolved in anhydrous THF (6 mL). The reaction was warmed to -20˚C and the solution of **3** was added dropwise over 30 min. The reaction was allowed to slowly warm to room temperature overnight, and acetic acid (0.5 mL) was added to yield an immediate blue color. The solvent was removed and the crude residue was dissolved in $CH_2Cl_2$ and washed with brine. The aqueous layer was extracted twice more with $CH_2Cl_2$, and the combined organic fractions were dried over $Na_2SO_4$. The solvent was evaporated, and the residue was redissolved in 5 mL of DCM and 5 mL of TFA. The solution was stirred for 2 h, then the solvent was evaporated and the crude product was purified by flash chromatography (0-10% MeOH in $CH_2Cl_2$) to yield a blue solid (79 mg, 49%) yield. $^{1}$H NMR (600 MHz, MeOD) δ 8.41 (s, 1H), 8.08 (dd, *J* = 7.9, 1.6 Hz, 1H), 7.26 – 7.21 (m, 3H), 6.62 (s, 2H), 4.36 (s, 2H), 3.60 (t, *J* = 5.6 Hz, 4H), 3.34 (s, 6H), 2.52 – 2.44 (m, 4H), 1.90 (p, *J* = 6.4 Hz, 4H), 0.57 (d, *J* = 6.7 Hz, 6H). $^{13}$C NMR (151 MHz, MeOD) δ 169.2, 165.8, 153.4, 148.4, 143.2, 141.5, 138.6, 132.6, 130.6, 129.3, 129.2, 128.3, 126.3, 121.2, 61.9, 53.5, 39.9, 28.2, 22.0, -1.0, -1.3. LRMS calc'd for $C_{31}H_{35}N_2O_3Si^+$ $[M]^+$ *m/z* 511.2411, found 511.6710.

**$HMSiR_{680}$-Me**: DIPEA (24 µL, 0.137 mmol, 4.7 eq) was added to a solution of $HMSiR_{THQ}$ (15 mg, 0.029 mmol) and TSTU (11 mg, 0.035 mmol, 1.2 eq) in anhydrous DMSO (0.129 mL) and the solution was stirred for 15 min. Methyl amine (33% in EtOH, 9 µL, 0.073 mmol, 2.5 eq) was added and the reaction was stirred at room temperature overnight. Excess base was removed by evaporation, and the resulting residue was purified by HPLC to yield the title compound as a blue TFA salt (11 mg, 58% yield). $^{1}$H NMR (600 MHz, MeOD) δ 8.19 (s, 1H), 7.88 (dd, *J* = 7.8, 1.7 Hz, 1H), 7.26 – 7.20 (m, 3H), 6.64 (s, 2H), 4.35 (s, 2H), 3.61 (t, *J* = 5.6 Hz, 4H), 3.34 (s, 6H), 2.99 (s, 3H), 2.52 – 2.43 (m, 4H), 1.91 (p, *J* = 6.3 Hz, 4H), 0.57 (d, *J* = 7.7 Hz, 6H). $^{13}$C NMR (151 MHz, MeOD) δ 170.2, 166.0, 153.4, 148.4, 141.8, 141.5, 138.7, 136.3, 130.6, 128.5, 126.9, 126.7, 126.2, 121.2, 62.1, 53.5, 39.9, 28.2, 27.0, 22.0, -1.0, -1.4. LRMS calc'd for $C_{32}H_{38}N_3O_2Si^+$ $[M]^+$ *m/z* 524.2728, found 524.6698.

### 3. NMR Spectra

Compound 2

$^{1}H$ NMR (500 MHz, $CDCl_3$)

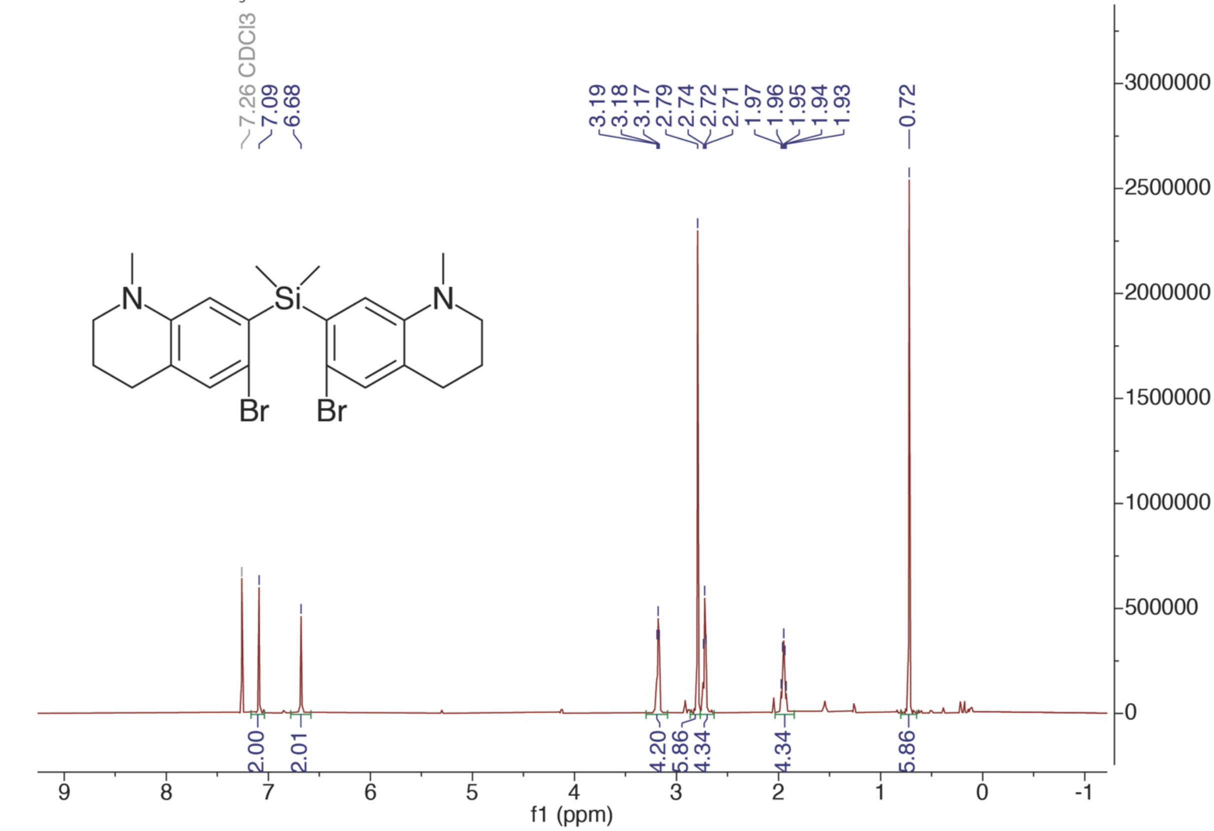

$^{13}C$ NMR (126 MHz, $CDCl_3$)

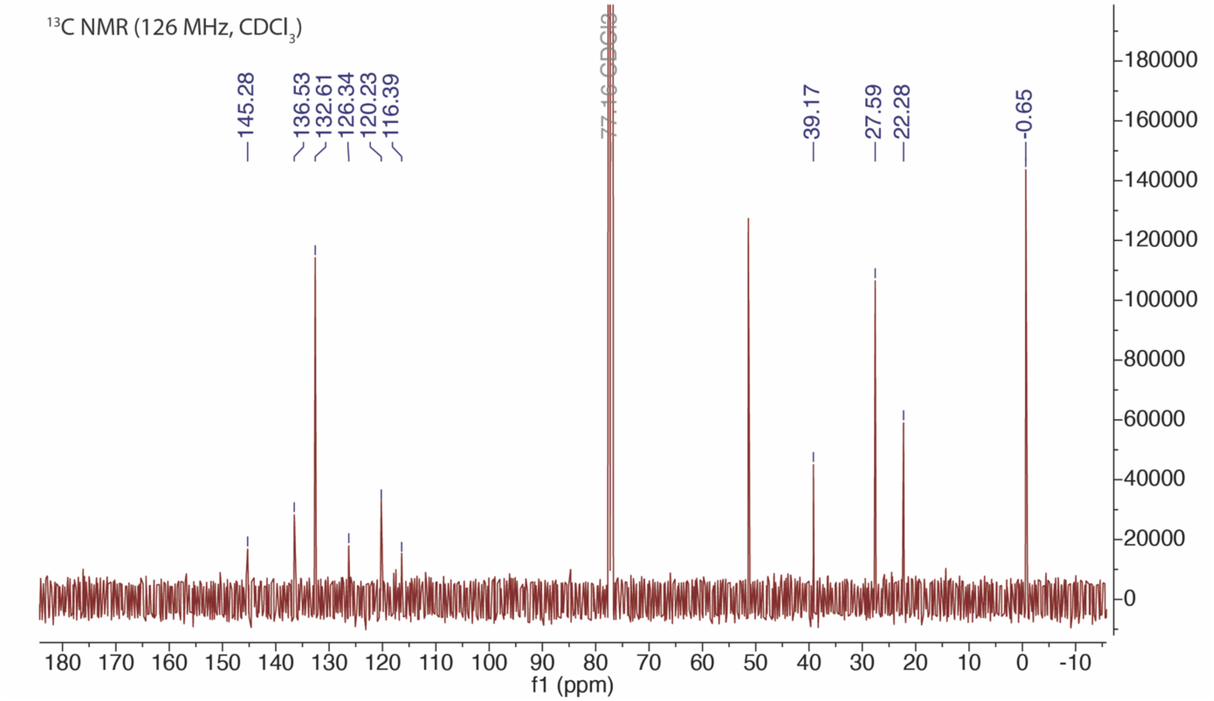

HMSiR$_{680}$

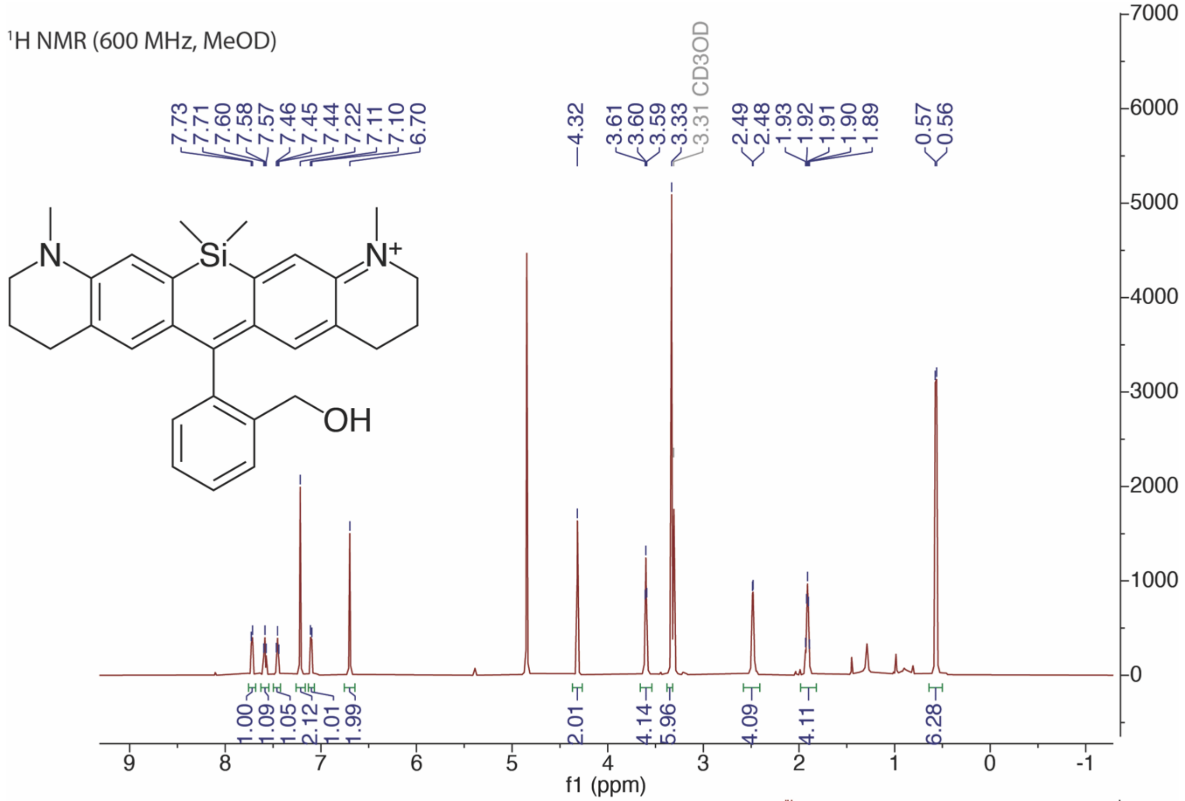

¹H NMR (600 MHz, MeOD)
7.73
7.71
7.60
7.58
7.57
7.46
7.45
7.44
7.22
7.11
7.10
6.70
4.32
3.61
3.60
3.59
3.33
3.31 CD3OD
2.49
2.48
1.93
1.92
1.91
1.90
1.89
0.57
0.56
N
Si
N+
OH
1.00
1.09
1.05
2.12
1.01
1.99
2.01
4.14
5.96
4.09
4.11
6.28
f1 (ppm)

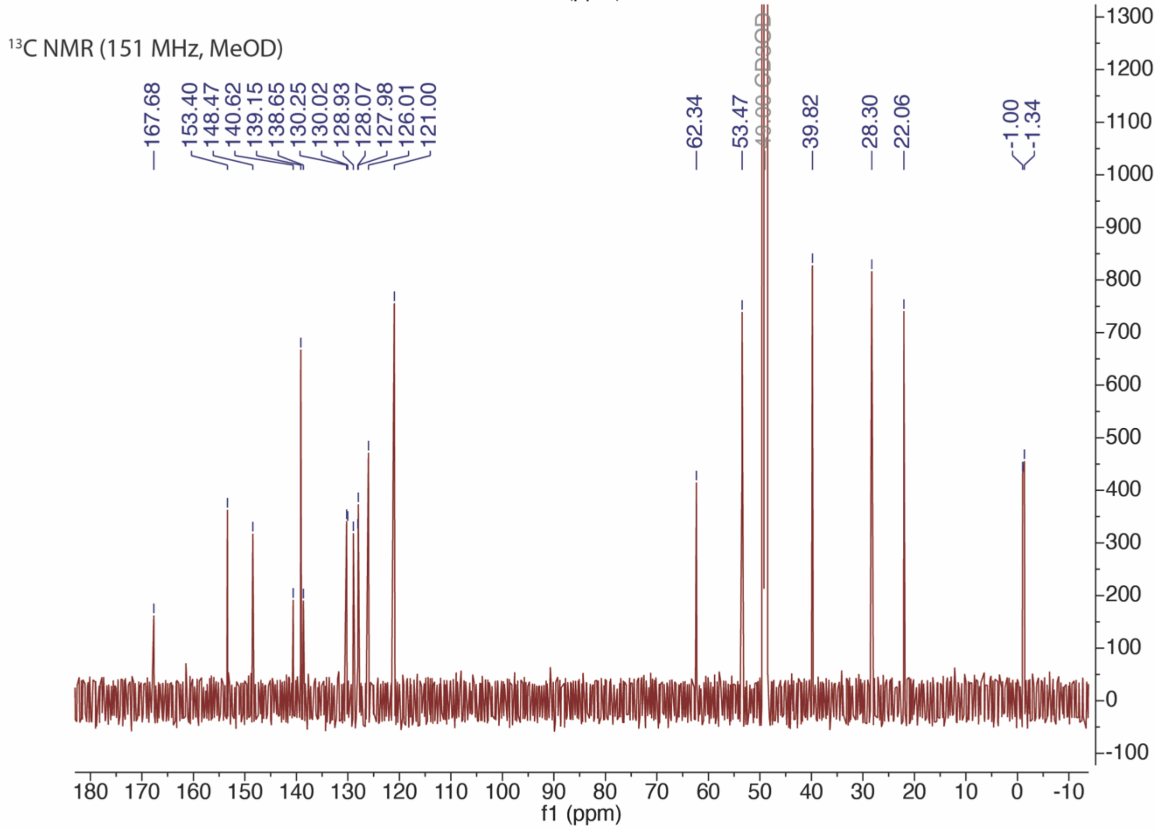

¹³C NMR (151 MHz, MeOD)
167.68
153.40
148.47
140.62
139.15
138.65
130.25
130.02
128.93
128.07
127.98
126.01
121.00
62.34
53.47
49.00 CD3OD
39.82
28.30
22.06
-1.00
-1.34
f1 (ppm)

HMSiR$_{THQ}$

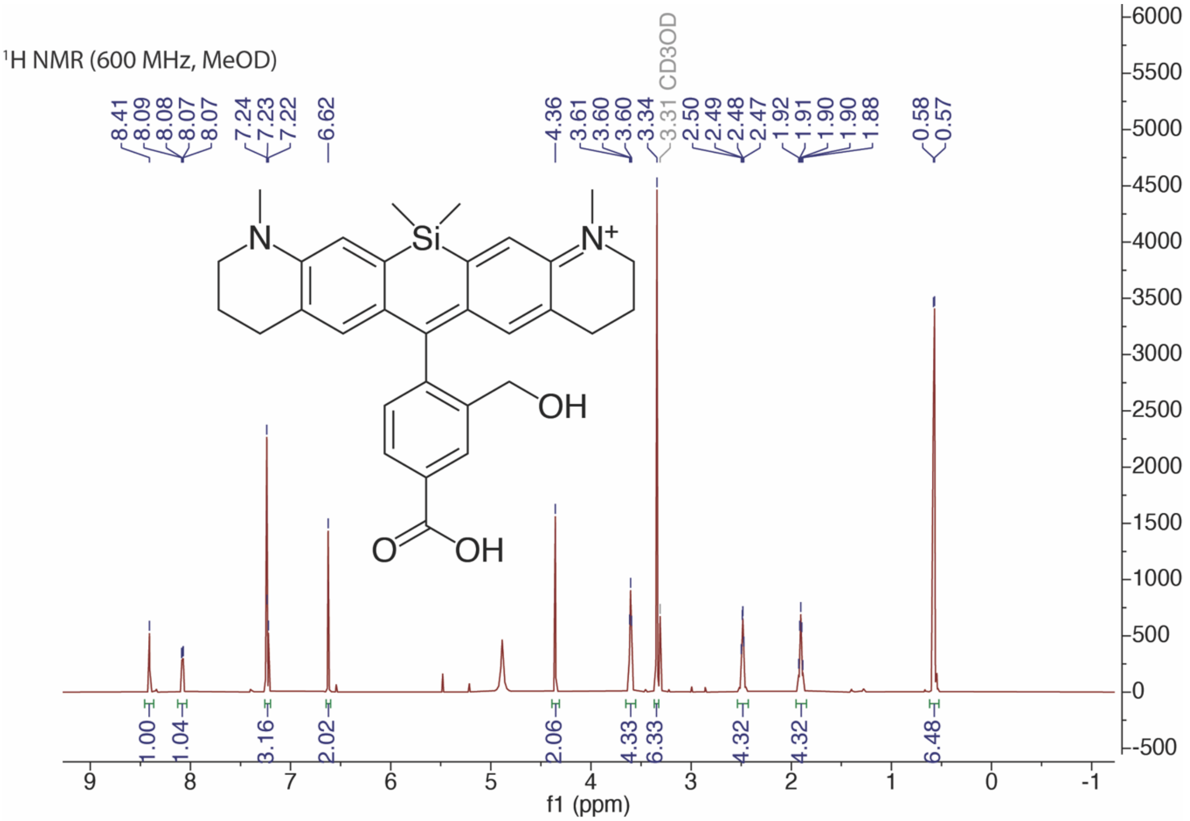

1H NMR (600 MHz, MeOD)
8.41
8.09
8.08
8.07
8.07
7.24
7.23
7.22
6.62
4.36
3.61
3.60
3.60
3.34
3.31 CD3OD
2.50
2.49
2.48
2.47
1.92
1.91
1.90
1.90
1.88
0.58
0.57
N
Si
N+
OH
O
OH
1.00
1.04
3.16
2.02
2.06
4.33
6.33
4.32
4.32
6.48
f1 (ppm)

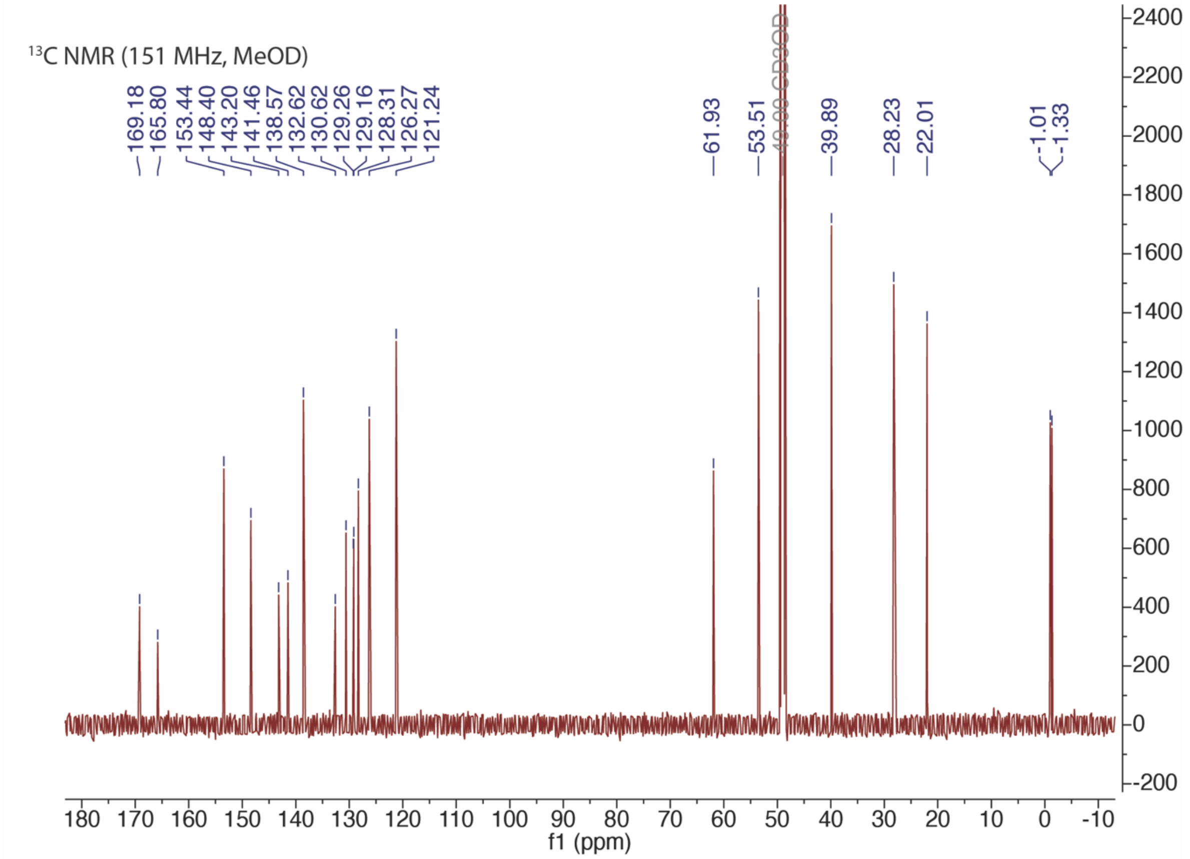

13C NMR (151 MHz, MeOD)
169.18
165.80
153.44
148.40
143.20
141.46
138.57
132.62
130.62
129.26
129.16
128.31
126.27
121.24
61.93
53.51
49.00 CD3OD
39.89
28.23
22.01
-1.01
-1.33
f1 (ppm)

HMSiR$_{680}$-Me

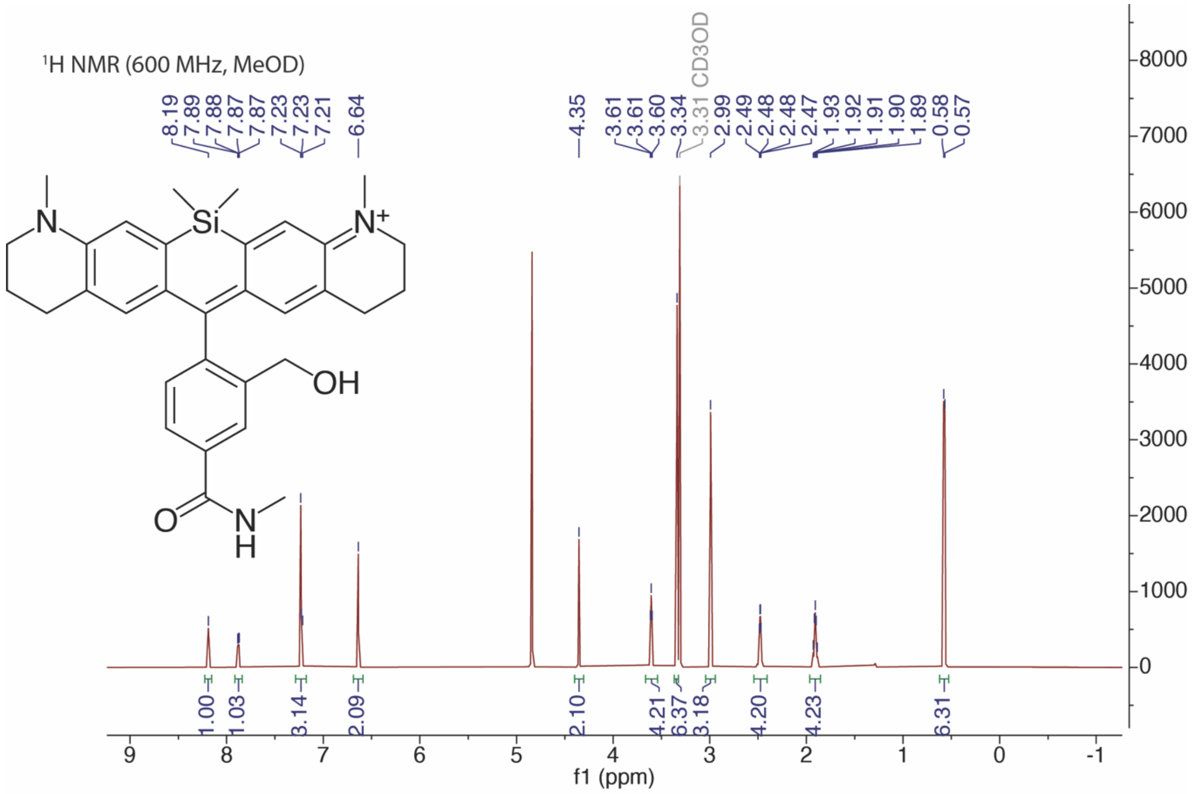

1H NMR (600 MHz, MeOD)
8.19
7.89
7.88
7.87
7.87
7.23
7.23
7.21
6.64
4.35
3.61
3.61
3.60
3.34
3.31 CD3OD
2.99
2.49
2.48
2.48
2.47
1.93
1.92
1.91
1.90
1.89
0.58
0.57
Si
N
N+
OH
O
N
H
1.00
1.03
3.14
2.09
2.10
4.21
6.37
3.18
4.20
4.23
6.31
f1 (ppm)

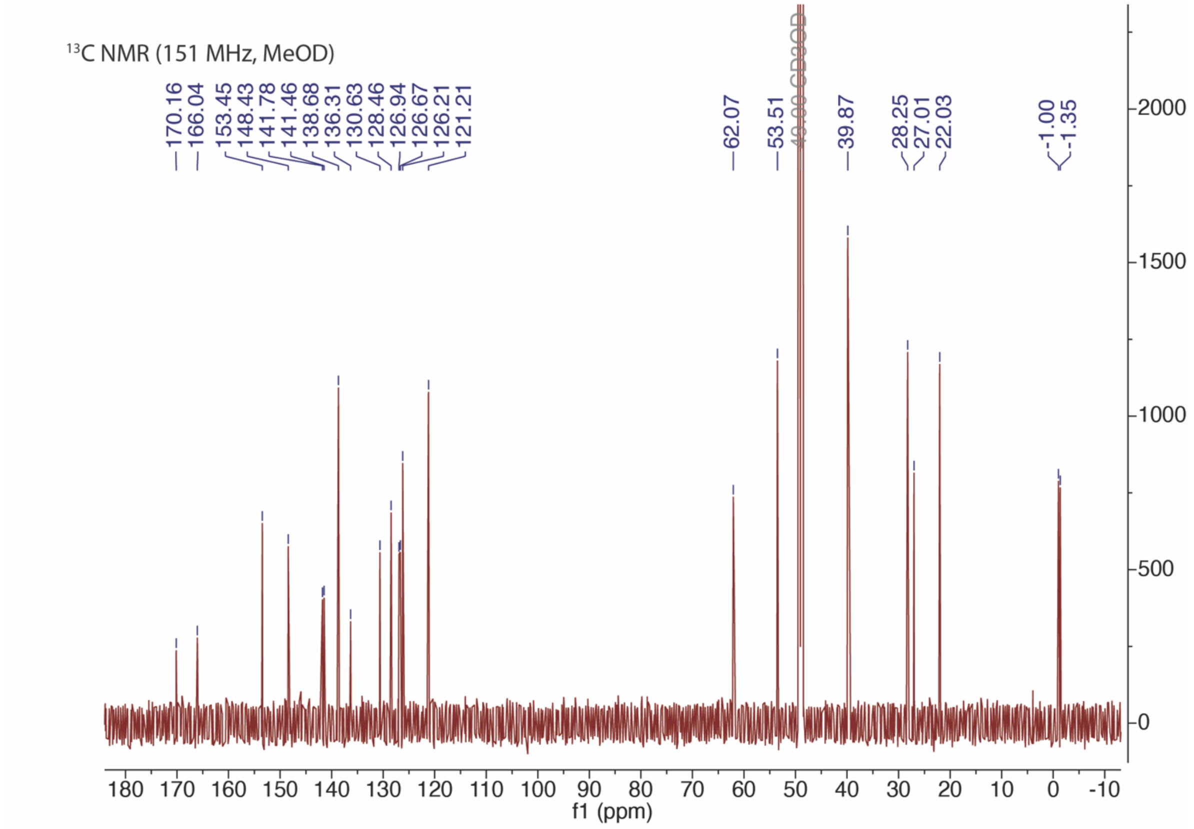

13C NMR (151 MHz, MeOD)
170.16
166.04
153.45
148.43
141.78
141.46
138.68
136.31
130.63
128.46
126.94
126.67
126.21
121.21
62.07
53.51
49.00 CD3OD
39.87
28.25
27.01
22.03
-1.00
-1.35
f1 (ppm)

### IV. References

(1) Stirling, D. R.; Swain-Bowden, M. J.; Lucas, A. M.; Carpenter, A. E.; Cimini, B. A.; Goodman, A. CellProfiler 4: Improvements in Speed, Utility and Usability. *BMC Bioinformatics* **2021**, *22* (1), 433. https://doi.org/10.1186/s12859-021-04344-9.

(2) Tinevez, J.-Y.; Perry, N.; Schindelin, J.; Hoopes, G. M.; Reynolds, G. D.; Laplantine, E.; Bednarek, S. Y.; Shorte, S. L.; Eliceiri, K. W. TrackMate: An Open and Extensible Platform for Single-Particle Tracking. *Image Process. Biol.* **2017**, *115*, 80–90. https://doi.org/10.1016/j.ymeth.2016.09.016.

(3) Grimm, J. B.; Brown, T. A.; Tkachuk, A. N.; Lavis, L. D. General Synthetic Method for Si-Fluoresceins and Si-Rhodamines. *ACS Cent. Sci.* **2017**, *3* (9), 975–985. https://doi.org/10.1021/acscentsci.7b00247.

(4) Tyson, J.; Hu, K.; Zheng, S.; Kidd, P.; Dadina, N.; Chu, L.; Toomre, D.; Bewersdorf, J.; Schepartz, A. Extremely Bright, Near-IR Emitting Spontaneously Blinking Fluorophores Enable Ratiometric Multicolor Nanoscopy in Live Cells. *ACS Cent. Sci.* **2021**, *7* (8), 1419–1426. https://doi.org/10.1021/acscentsci.1c00670.

(5) Das, B.; Venkateswarlu, K.; Majhi, A.; Siddaiah, V.; Reddy, K. R. A Facile Nuclear Bromination of Phenols and Anilines Using NBS in the Presence of Ammonium Acetate as a Catalyst. *J. Mol. Catal. Chem.* **2007**, *267* (1), 30–33. https://doi.org/10.1016/j.molcata.2006.11.002.
